# The Ketosynthase Domain Constrains the Design of Polyketide Synthases

**DOI:** 10.1101/2020.01.23.916668

**Authors:** Maja Klaus, Lynn Buyachuihan, Martin Grininger

## Abstract

Modular polyketide synthases (PKSs) produce complex, bioactive secondary metabolites in assembly line-like multistep reactions. Longstanding efforts to produce novel, biologically active compounds by recombining intact modules to new modular PKSs have mostly resulted in poorly active chimeras and decreased product yields. Recent findings demonstrate that the low efficiencies of modular chimeric PKSs also result from rate limitations in the transfer of the growing polyketide chain across the non-cognate module:module interface and further processing of the non-native polyketide substrate by the ketosynthase (KS) domain. In this study, we aim at disclosing and understanding the low efficiency of chimeric modular PKSs and at establishing guidelines for modular PKSs engineering. To do so, we work with a bimodular PKS testbed and systematically vary substrate specificity, substrate identity, and domain:domain interfaces of the KS involved reactions. We observe that KS domains employed in our chimeric bimodular PKSs are bottlenecks with regards to both substrate specificity as well as interaction with the ACP. Overall, our systematic study can explain in quantitative terms why early oversimplified engineering strategies based on the plain shuffling of modules mostly failed and why more recent approaches show improved success rates. We moreover identify two mutations of the KS domain that significantly increased turnover rates in chimeric systems and interpret this finding in mechanistic detail.

## INTRODUCTION

Multimodular polyketide synthases (PKSs) are large megasynthases, which are responsible for the production of a variety of pharmaceutically important compounds such as antibiotics, anti-fungal and anti-cholesterol agents (Figure 1A).^1^ The modularity of PKSs has put them at the forefront of engineering efforts to achieve programmable access to novel compounds with new bioactivities. However, to date most of the engineered systems do not reach biotechnological relevant stages, because the turnover numbers and thus product yields are often too low.^2–4^ Despite considerable efforts in the last 30 years, no generalizable rules for the generation of efficient chimeric PKSs exist today. For example, although the importance of protein-protein interactions e.g. between docking domains^5–7^ or between the acyl carrier protein (ACP) and the ketosynthase (KS) at different stages in the catalytic cycle has been recognized (Figure 1B),^4,8–10^ most strategies to modulate and adapt those interactions in chimeric modular PKSs have failed so far.^11–14^

**Figure 1.**
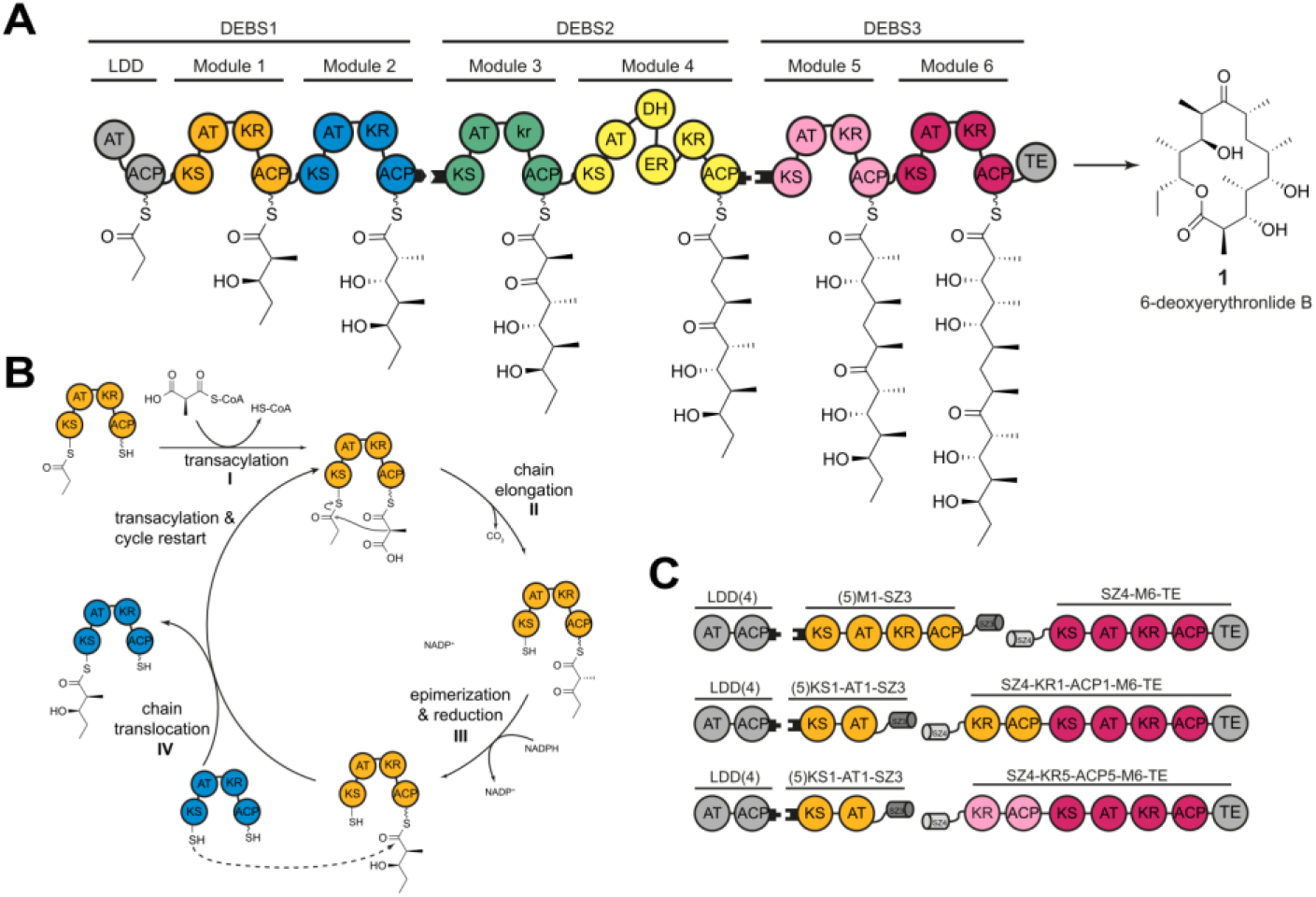
Schematic architecture of DEBS, its catalytic cycle and accessible interfaces for engineering approaches. (A) Architecture of DEBS. The three polypeptides (DEBS1-3), the encoded modules (M1-M6), as well as the loading didomain (LDD), the thioesterase domain (TE), and the final product (**1**) are depicted. Polyketide intermediates are shown as attached to the respective acyl carrier protein (ACP). Black tabs depict docking domains. Domain annotations are as follows: LDD-loading didomain, AT-acyltransferase, KS-ketosynthase, ACP-acyl carrier protein, KR-ketoreductase, DH-dehydratase, ER-enoylreductase, TE-thioesterase. Module 3 has a ketoreductase-like domain (denoted in lowercase) that lacks NADPH-dependent oxidoreductase activity but harbors C2 epimerase activity.^27,28^ (B) Catalytic cycle on the example of M1 and M2. (I) AT-catalyzed transacylation using (2*S*)-methylmalonyl-CoA, followed by KS-mediated decarboxylative Claisen condensation (chain elongation, II) to give the (2*R*)-2-methyl-3-ketoacyl-ACP.^29^ In M1 the KR catalyzes the epimerization of the C2 methyl group, followed by diastereospecific reduction (III) to give the (2 *S*,3*R*)-2-methyl-3-hydroxydiketide product. The diketide is translocated to the KS domain of the downstream, substrate-accepting module (chain translocation, IV) from which the next round of reactions can start. In DEBS only KR1 and KR3 catalyze epimerization of the C2 methyl group.^30^ (C) Subset of bimodular chimeric PKSs used previously.^22^ SYNZIP domains (SZ3 and SZ4) assisted in construct design. Black tabs represent DEBS-derived docking domains (module numbers in brackets) installed at the protein termini to mediate weak interactions.

The 6-deoxyerythronolide B synthase (DEBS) is the prototypical example of this enzyme class and has served as a platform to study the structure, mechanism, and engineering potential of these synthases (Figure 1A).^15^ In the past, numerous studies on DEBS have shed light on substrate tolerance,^6,16^ protein-protein interaction specificities,^5,9,17,18^ and structural characteristics^19–21^ with broad relevance for the family of modular PKSs. Recently, we analyzed a library of bimodular chimeric PKSs, with modules mainly deriving from DEBS, to compare different chimeric interfaces (Figure 1C).^22^ From our analysis, it became apparent that some systems in which native domain:domain interfaces have been preserved across chimeric module boundaries (from the ACP of the upstream module to the KS of the downstream module) still suffer a decrease in turnover rates. We have interpreted those results as kinetic penalties in turnover rates arising from the narrow specificity of the domains dealing with the non-cognate substrates.^22^

There are multiple options for a downstream, substrate-accepting module to be rate-limiting when processing a non-cognate substrate. In case of our bimodular PKS testbed (Figure 1C, bottom), three reactions can in principle account for substrate specificity of the substrate-accepting module (M6-TE) and cause the aforementioned phenomenon; (i) the KS6-catalyzed condensation reaction, (ii) the following NADPH-dependent reduction of the β-ketoester by KR6, or (iii) the TE-catalyzed lactonization to the triketide product. While no information is available on the steady-state kinetics of the individual reaction steps within M6-TE, previous analysis of the hydrolysis rate of two enantiomeric diketides by the TE domain,^23^ comparison of the reduction rates of two KR domains^24^ and kinetic analysis of the KS domain of DEBS M2^16,25^ have led to the conclusion that the KS-catalyzed condensation likely limits the rate of a native module.^15,25^ Identification of substrate specificity of the KS domain as a bottleneck in PKS turnover is further supported by a recent mutagenesis study on DEBS KS3 in which single point binding site mutagenesis resulted in significantly broadened substrate specificities and higher turnover rates.^26^ Based on the observation that the KS domains of the mycolactone synthase (MYCL) share a >97% sequence identity, despite accepting substrates of different length and chemical modification, DEBS KS3 residues were replaced with their counterparts from MYCL, resulting in one particular point mutant (A154W_KS3_) with significantly increased promiscuity compared to wild-type KS3.^26^

Following the argument that the KS domain of the substrate-accepting module is throttling turnover rates in chimeric PKSs when interacting with a non-cognate ACP or processing a non-cognate substrate, we started a study to more precisely understand KS in these properties. Similarly, we aimed at probing to which extend KS binding site engineering can improve the turnover of noncognate substrates. For addressing these questions, we were building on our prior experience with the bimodular DEBS *in vitro* testbed. Phylogenetically, DEBS KS domains are situated in the same clade as KS domains from other Actinobacterial modular type I PKS, which means they are closely evolutionarily related and likely similar in sequence.^31–34^ As such, the DEBS system is not only the most widely studied modular PKS, but studies on its KS domains are also representative for type I modular PKS KS domains.

In our chimeric bimodular testbed, we varied essentially three parameters, all related to the KS of the substrate-accepting module; (i) substrate specificity, (ii) substrate identity, and (iii) domain:domain interfaces. Substrate specificity was varied by designing multipoint mutations under guidance of the FuncLib program and consideration of Rosetta energies to preserve protein stability.^35^ Mutated constructs showed changed efficiencies, i.e. increased or released kinetic penalties imposed by the substrate, demonstrating that substrate specificity of the KS is a rate-determining factor in chimeric PKSs. For some multipoint mutants, that were particularly powerful in increasing turnover rates on the chimeric systems, we give information at mechanistic level. Finally, we altered the substrate-donating module in the chimeric constructs to also vary protein-protein interactions and substrate identity. In conclusion, our data demonstrates that the condensation reaction in our chimeric modular PKSs, which comprises the translocation of the growing polyketide chain from one module to the other (transacylation, ping step) and the elongation by a C2-unit (decarboxylative condensation, pong step), is constrained by domain-domain interactions and substrate specificity of the KS domain. We believe our data can explain why engineering approaches based on the simple shuffling of modules have often failed in the past. In addition, we show that multipoint mutagenesis within the KS binding site can increase the turnover rate of chimeric PKSs.

## RESULTS AND DISCUSSION

### The Bimodular DEBS Testbed

We aimed at probing KS properties *in vitro* to avoid uncontrolled interference with physiological processes of a host cell and to allow a simple and direct read-out of enzymatic properties. Just a few modular PKSs are available for *in vitro* analysis to date, i.e. the venemycin synthase,^36^ the phoslactomycin synthase,^37^ the nocardiosis-associated polyketide synthase,^38^ and the DEBS synthase^39^. Since most of these systems were just recently reported, solely the DEBS PKS has been thoroughly established for *in vitro* studies during the last years. In this study, we were working with two DEBS-based chimeric bimodular systems, differing in their substrate-accepting module; i.e., (i) LDD(4)+(5)M1-SZ3+SZ4-M3-TE and (ii) LDD(4)+(5)M1-SZ3+SZ4-M6-TE (Figure 2A). Both bimodular systems harbor a non-cognate interface (ACP1:KS3 or ACP1:KS6) and challenge the KS of the substrate-accepting module (KS3 or KS6) with the non-cognate DEBS-derived natural diketide (NDK, (2*S*,3*R*)-2-methyl-3-hydroxydiketide (Figure 2A). Note that docking domains (indicated as (4)+(5))^39^ and SYNZIP domains (indicated as SZ3+SZ4)^40^,^41^ were employed at the protein interfaces to mediate non-covalent interactions between separately purified modules (Figure 2A). The choice of M3 and M6 was guided by their differing turnover rates with M1^22^ and the previously observed substrate restrictions.^16^ Further, X-ray structural data is available for KS3-AT3,^42,43^ which allows mapping structure-function relationships.

**Figure 2.**
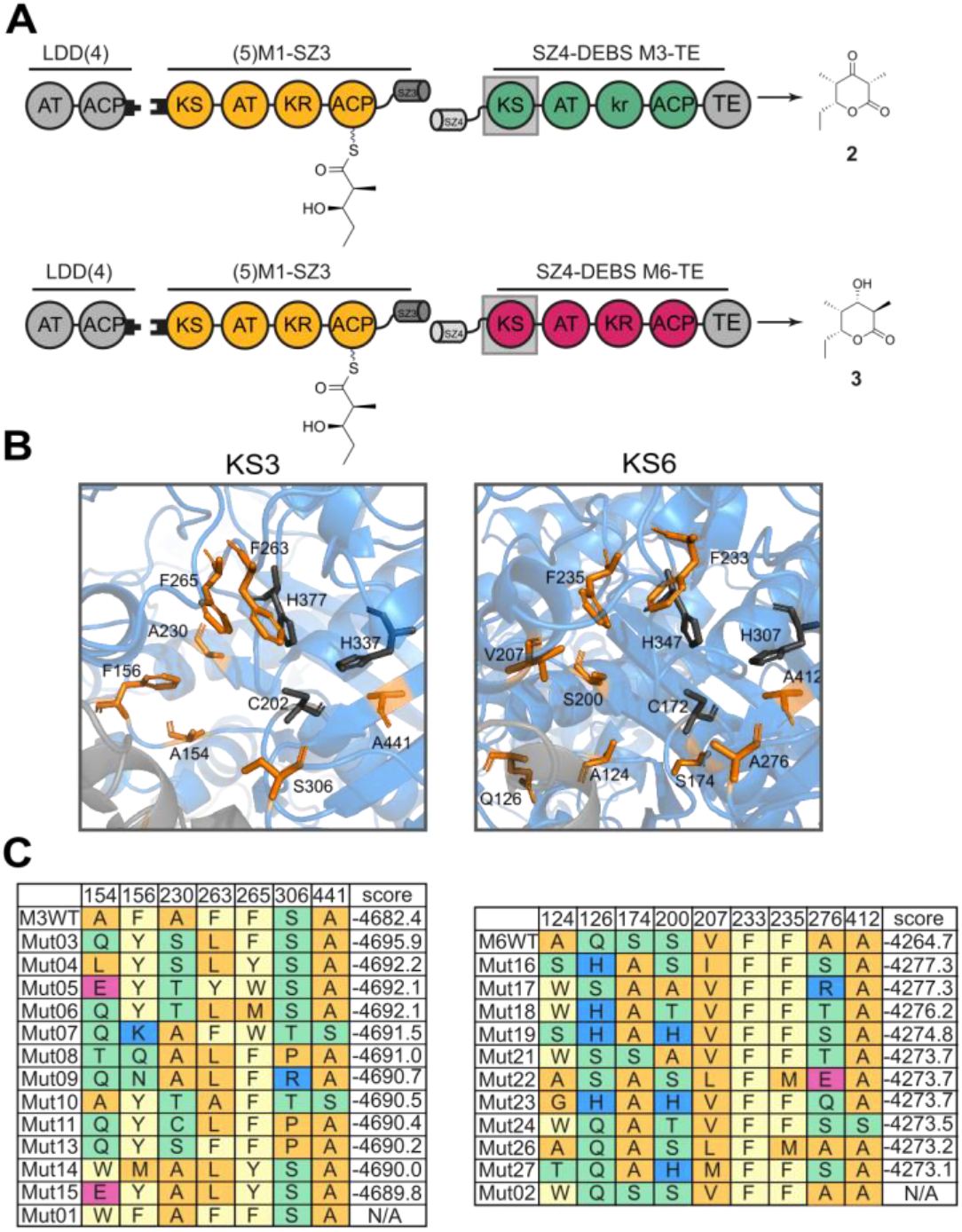
KS mutagenesis - Setup of bimodular chimeric PKSs, structural organization of KS binding sites as well as an overview of KS3 and KS6 mutants. (A) Bimodular chimeric PKSs using LDD(4), (5)M1-SZ3, and SZ4-M3-TE or SZ4-M6-TE as the substrate-accepting module. Matching DEBS-derived docking domains (black tabs; (4)+(5)) or matching SYNZIP domains (SZ3+SZ4) are installed at the protein termini to mediate non-covalent interactions.^22^ The substrate-donating ACP1 harbors the (2*S*,3*R*)-2-methyl-3-hydroxydiketide (natural diketide, NDK) as the incoming substrate for wild-type and mutant KS domains. Depending on the substrate-accepting module either unreduced triketide lactone (TKL) **2** or reduced TKL **3** are the predicted products. (B) Structural organization of the binding sites of KS3 and KS6. The catalytic triad is depicted as gray sticks. Residues chosen for mutagenesis within a 12 Å-range of the active site cysteine C202_KS3_ or C172_KS6_ are shown as orange sticks. (C). Selection of KS3 and KS6 mutants sorted by Rosetta Energy in REU. Amino acids are colored according to their side chain properties. Mutants were selected to exhibit great variability and include some of the residues found in the highly conserved KS domains of MYCL (Figures S1&S2). In addition, single point mutants Mut01 (A154W) and Mut02 (A124W) were created based on previous results.^26^ Rosetta scores were calculated based on the KS-AT didomain. N/A: not applicable. The sequence space for each position is listed in Table S1.

### Design and Generation of KS Multipoint Mutants

To explore KS specificity in constraining turnover rates of chimeric PKSs, particularly aiming at finding ways to release these constraints to increase chimeric PKS efficiency, we first modulated the binding site of the KS domain of the substrate-accepting module of our bimodular PKSs (Figure 2B). We hypothesized that multipoint mutations within KS binding sites will have a higher impact on substrate specificity than single point mutations. Based on a set of manually selected residues, we used the FuncLib approach for generating a variety of diverse multipoint mutants (Figure 2C).^35^ FuncLib uses phylogenetic information to reveal possible, coupled mutations within a protein by working on two levels: First, it uses phylogenetic information to suggest amino acid exchanges at selected positions, and, second, it ranks the mutated proteins by calculating their stability via the Rosetta program suite. In this process, mutations that are predicted to destabilize the protein fold are discarded. The FuncLib approach does not target individual substrate specificities but delivers a set of stable variants that can then be screened for the specificities and activities of interest.

In applying FuncLib to our bimodular systems, we used X-ray structural data for KS3-AT3,^42,43^ to calculate stable multipoint mutations for module M3 (construct SZ4-M3-TE). For M6 (construct SZ4-M6-TE), a homology model of KS6-AT6 was generated on the basis of a sequence similarity of ~56% to the structurally solved didomain KS5-AT5.^19^ For both modules, we initially spotted residues for KS binding site mutagenesis within a shell around the active cysteines (Figures S1&S2). From this set of residues, we determined the non-conserved ones as suited for mutagenesis, as they are likely involved in generating substrate specificity. With this approach, a total of 7 residues for KS3 and 9 residues for KS6 were received (Figure 2C). FuncLib generated a sequence space of 152,826 designs for KS3-AT3 and 285,822 designs for KS6-AT6 with a large variability in mutated residues (Table S1). Rosetta calculation eventually revealed that from the initial designs 1,011 designs for KS3-AT3 and 3,683 designs for KS6-AT6 result in more stable proteins than the respective wild-type proteins. From the 50 highest ranked designs, we chose 12 for KS3-AT3 and 10 for KS6-AT6 for subsequent analysis, based on their large variability in mutated residues (Figure 2C). All designs harbored a minimum of three and a maximum of five mutations. In addition, the previously reported single point mutants A154W_KS3_ and A124W_KS6_ were also generated as benchmark.^26^ The mutations are predicted to change the geometry of the respective binding sites, potentially leading to different properties in substrate binding and thus activity (Figures S3&S4).

To evaluate the performance of the different KS designs under PKS turnover conditions, all KS mutations were introduced in the background of the full-length modules SZ4-M3-TE and SZ4-M6-TE. All mutant proteins exhibited similar purification behavior and purity (Figures S5A&B), with yields comparable or higher than the wild-type modules (between 5 - 17 mg per liter *E. coli* culture, Table S2). Protein oligomerization, as measured by SEC, showed that besides Mut07 and Mut10 all mutant proteins eluted in a single peak indicative of a homodimeric protein fold (Figures S6&S7). In good agreement with these results, thermal shift assays of each mutant indicated no significant difference in thermal stability between wild-type and mutant modules (Table S3). In confirming high protein quality, we were able to have an undisturbed view on the impact of KS binding site design on the turnover rates of chimeric PKSs.

### Analyzing the Turnover Rates of KS Multipoint Mutants

FuncLib-assisted design of binding sites cannot be directed to increase specificity for substrates and ligands.^35^ In analyzing KS domains as putative bottlenecks in chimeric PKSs, we thus primarily aimed at altered and not *per se* at increased turnover rates. We first assessed the turnover rates in the bimodular chimeric PKS LDD(4)+(5)M1-SZ3+SZ4-M3-TE (Figure 3A). As observed before, the chimeric system with wild-type SZ4-M3-TE exhibited a >30-fold lower activity than the reference system with SZ4-M2-TE (Figure 3A, gray bar).^22^ Notably, compared to wild-type M3, mutated M3 showed changed turnover rates, revealing KS3 as constraint in the chimeric bimodular PKSs. The single point mutant Mut01 (A154W_KS3_), following the design of Murphy *et al*.,^26^ had a roughly 2-fold and the multipoint mutants Mut04, Mut08, Mut09, Mut11 and Mut13 a roughly 2.5-fold increase in turnover rates. In good agreement with the turnover rates, product analysis after overnight incubation confirmed the presence of the expected unreduced TKL **2** in all systems with measurable turnover (Figure S8).

**Figure 3.**
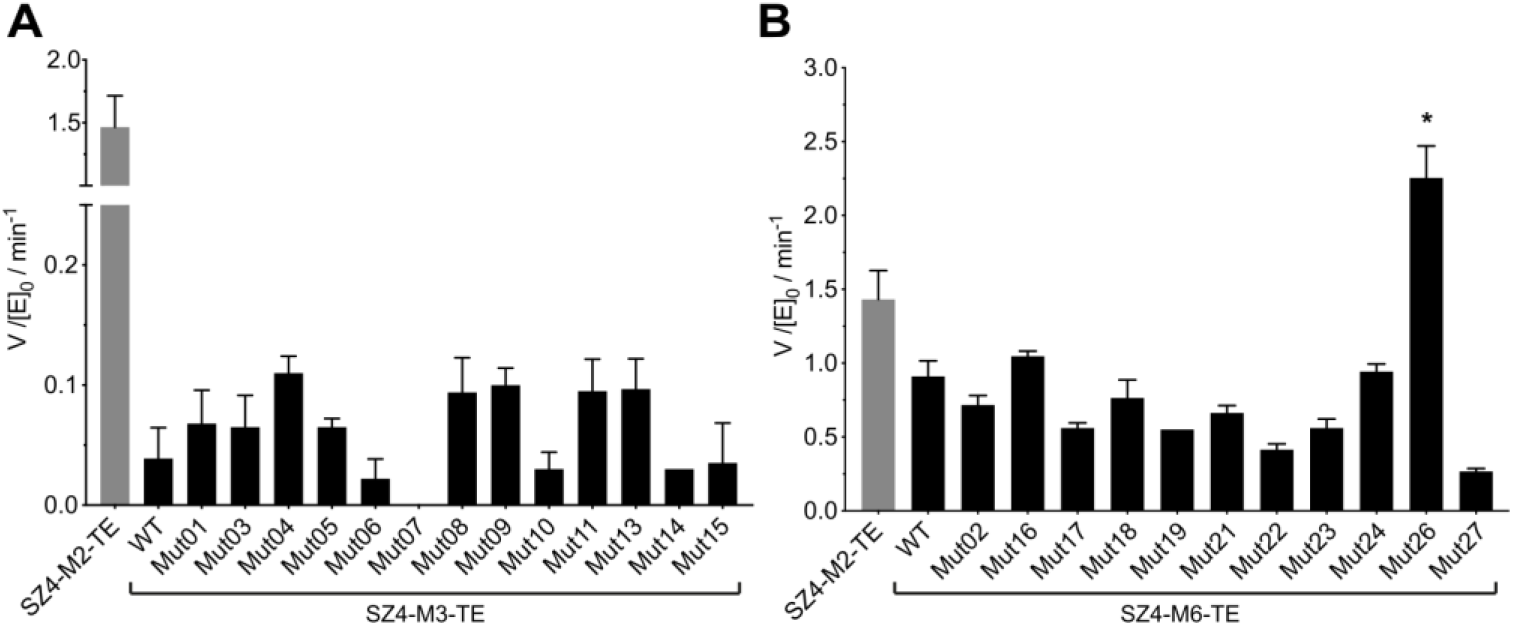
Turnover rates of bimodular chimeric PKSs using wild-type or mutant M3-TE (A) and M6-TE (B), respectively. Bimodular PKSs consisted of LDD(4), (5)M1-SZ3, and SZ4-M3-TE/SZ4-M6-TE (wild-type (WT) or one of their mutants). For both bimodular PKSs, the reference system consisted of LDD(4), (5)M1-SZ3, and SZ4-M2-TE (gray bar). All initial rate data was obtained at 4 μM enzyme concentration and non-limiting concentrations of propionyl-CoA, methylmalonyl-CoA, and NADPH. Measurements were performed in technical triplicate. Note that for further confidence, the turnover rate of Mut26 is displayed as an average of three independently purified protein samples (biological triplicate) measured in triplicate.

The second bimodular chimeric PKSs tested in this study, LDD(4)+(5)M1-SZ3+SZ4-M6-TE, responded similar to mutations in the KS (KS6) of the substrate-accepting module (M6) (Figure 3B). All mutants except mutant Mut26 showed no remarkable improvement or decreased turnover rates. Interestingly, Mut26 did not only show significantly increased turnover compared to wild-type M6 (~2.5-fold improvement) but performed even better than the reference system M2-TE (1.5-fold improvement, Figure 3B). Note that for all substrate-accepting modules shown in Figure 3B, a mixture of the expected reduced TKL **3** and unreduced TKL **5** was obtained from the reaction mixture after a reaction time of only 10 min (Table S4, Figure S9). Similar data has been reported before for DEBS M6-TE.^44^ As the turnover rates were measured by correlating the reduction rates of the KR domains (NADPH consumption) with product formation and assuming a 100% conversion to reduced TKL,^39^ the absolute turnover numbers presented in Figure 3B would all be higher if corrected by the amount of produced unreduced TKL. Yet, as the ratios of reduced to unreduced TKL are similar in all M6 samples (Table S4), the relative ratios of turnover rates are still captured in Figure 3B.

### Analyzing the Effect of Domain Swaps on the Turnover Rates of Chimeric PKSs

By employing (5)M1-SZ3 as the substrate-donating module in the tested chimeric PKSs, our analysis gave insight into the effect of each mutant when presented with the non-cognate ACP1 bearing NDK (Figure 2A). With the aim of additionally analyzing the contribution of ACP:KS recognition and substrate-KS interaction on the turnover of KS mutants, we generated alternative M1 substrate-donating modules (Figure 4A). All substrate-donating proteins, presented in the following, were purified in similar quantities and were pure as judged by SEC and SDS-PAGE (Table S4, Figures S10&S5C). Eventually, the new modules were assembled with a subset of substrate-accepting multipoint mutants to generated novel chimeric PKSs. While we focused on the M3-based mutants, Mut01, Mut08, Mut11, Mut13 and Mut15, we also tested the Mut26-mutant of the M6 module due to its unexpected high turnover rates (Figure 4A).

**Figure 4.**
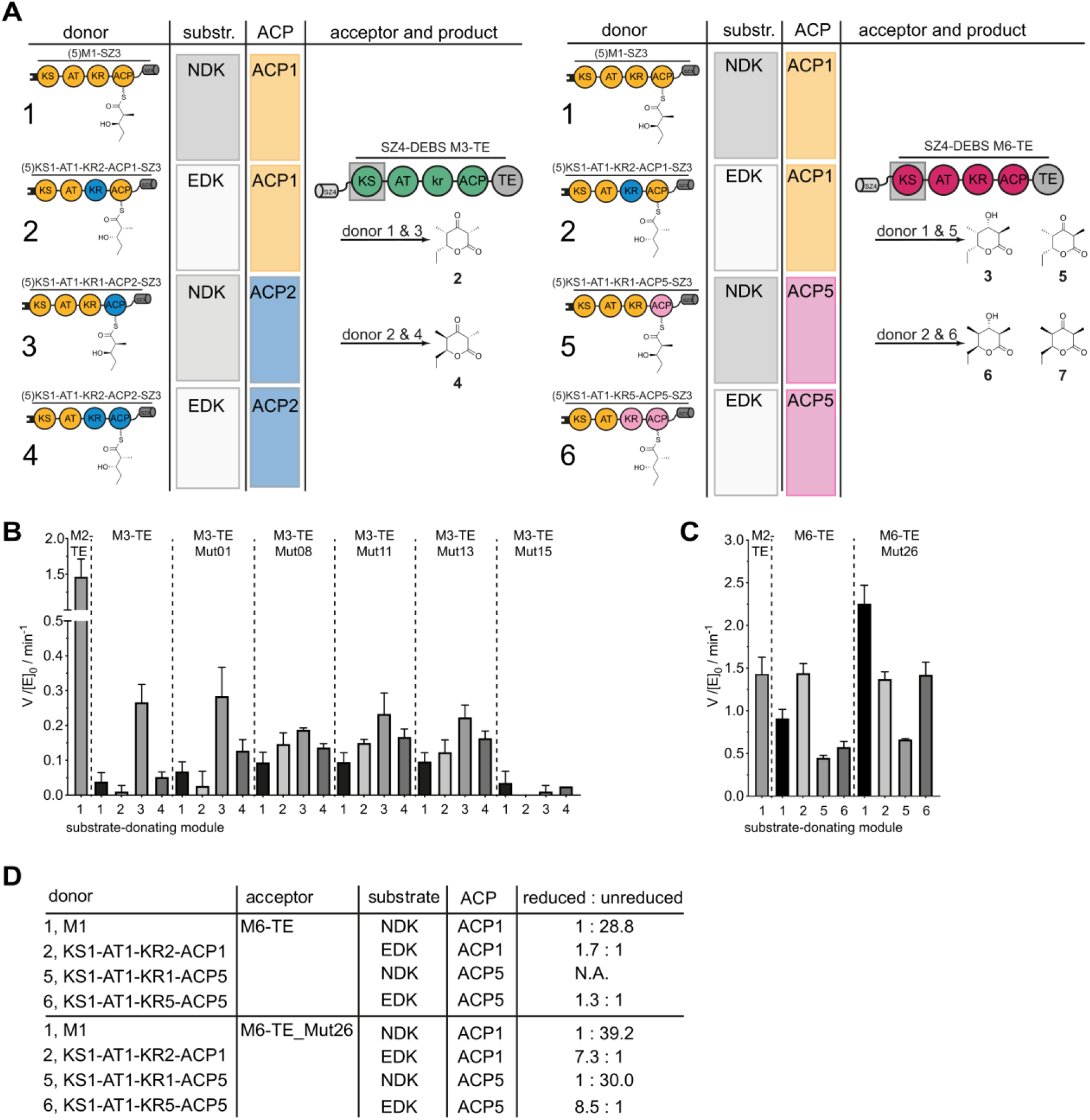
Turnover rates of bimodular chimeric PKSs using M3-TE, M6-TE and different substratedonating modules. (A) The substrate-donating modules used to generate bimodular PKSs. Substrates, the employed ACPs, and generated products are indicated. Turnover rates of bimodular chimeric PKS using M3-TE (B) or M6-TE (C) as substrate-accepting modules. All bimodular PKSs consisted of LDD(4), followed by a substrate-donating module as indicated in (A), and SZ4-M3-TE, SZ4-M6-TE or one of their mutants. The reference system consisted of LDD(4), (5)M1-SZ3, and SZ4-M2-TE (gray bar). All initial rate data was obtained at 4 μM enzyme concentration and non-limiting concentrations of propionyl-CoA, methylmalonyl-CoA, and NADPH. Measurements were performed in technical triplicate. The expected TKLs **2-7** were detected in all samples after overnight incubation (Figures S8, S9&S11). (D) Products of bimodular PKSs using M6-TE or Mut26. Different ratios of reduced to unreduced species were produced with different donors, yet wild-type M6 and Mut26 show the same trends. Calculation based on peak areas shown in Table S5. N. A., not applicable.

In summarizing this data, the engineering of domain swaps is outlined first, and then turnover rates are reported. Three types of domains swaps were generated (Figure 4A): (i) Since KR domains are responsible for installing the stereogenic centers at the α- and β-position of the polyketide, the exchange of KR1 by KR2 allowed generating the DEBS-derived enantiomeric diketide (EDK, (2*R*,3*S*)-2-methyl-3-hydroxydiketide) on ACP1 *in situ.^4^* With this construct, we were able probing the impact of substrate identity on the turnover rates of KS-mutated chimeras. Note that the KR-swap causes a non-cognate KR2-ACP1 interaction within M1. Based on the observations that KR domains can efficiently turn over with a variety of ACPs,^45^ we did not expect this interface to be rate-limiting (Figure 4A, donor 2). (ii) As a next step, we swapped ACP1 of the substrate-donating M1 with ACP2 and ACP5, the native upstream ACPs of M3/M6. In doing so, we generated substrate-donating modules that carry NDK on the respective ACP and promote translocation via a cognate ACP:KS interface (Figure 4A, donor 3 and 5). As for the KR exchange, also the swap of ACP creates noncognate interfaces in M1. As elongation at a chimeric KS:ACP interface in a fusion protein is not rate-limiting,^22^ we again did not expect to install a rate-limiting kinetic penalty. (iii) As a last engineering step, both KR1 and ACP1 were exchanged by the native upstream domains of M3/M6, resulting in the formation of EDK on either ACP2 (Figure 4A, donor 4) or ACP5 (Figure 4A, donor 6).

The KR-swaps were designed to highlight the sensitivity of the KS within the accepting module for the enantiomeric substrates NDK and EDK. Changes in turnover rates turned out to be rather small for the M3 containing bimodular PKS (Figure 4B). The ACP swaps proved to be more effective in changing turnover rates. Donor 3, that when coupled to the substrate-accepting module M3 reconstitutes the native ACP2-KS3 interaction and presents KS3 with the NDK substrate, improved rates with wild-type and all mutated acceptors except Mut15 (Figure 4B). Donor 4 appeared as the second-best substrate-donating module for M3, again employing ACP2, but displaying the enantiomeric EDK (Figure 4B). The comparison of different donors for the M3 substrate-accepting module, particularly for wild-type M3 and Mut01, highlights the importance of protein-protein interactions at the ACP:KS interface in our chimeric bimodular PKS testbed. As another trend in our data, wild-type M3 and the single point mutant Mut01 showed a strong substrate preference for NDK over EDK, while multipoint mutants tended to broaden the substrate tolerance of KS3 and release the ACP preference.

The analysis of bimodular chimeric PKSs with wild-type M6 and Mut26 as substrate-accepting modules turned out to be more difficult (Figure 4C). As mentioned above, all bimodular PKSs using wild-type M6 or Mut26 produced a mixture of reduced and unreduced TKLs (Table S4, Figure S9), which hampers kinetic analysis of the bimodular PKSs and, specifically, underestimates the actual number of turnovers. Intriguingly, we faced an additional effect, further complicating interpretation of data. While we received similar ratios of reduced to unreduced species for wild-type and mutated M6 modules (Table S4, Figure S9), the relative ratios varied with the employed substrate-donating module (Figure 4D, Table S5). Since only the KR domain within the substrate-accepting module defines the ratio of reduced to unreduced species, this phenomenon has to be explained by an impact of the incoming substrate on KR6 activity. Likely, a pronounced substrate specificity of KR6, observed by others before,^46^ accounts for the impact of the substrate-donating module, and causes the more efficient reduction of the triketide originating from EDK compared to the triketide from NDK (Figure 4D). Overall, the analysis of the chimeric bimodular PKSs containing wildtype and Mut26-mutated modules did not reveal an unambiguous correlation between turnover rates in chimeric PKSs and involved protein-protein interactions or substrate-KS interactions, respectively. However, as observed for the M3-containing bimodular PKSs, the multipoint mutations broadened the substrate tolerance of KS6 and exhibited a released ACP preference.

### Mechanistic Insight Into Chimeric PKS

In probing catalytic efficiencies of 24 KS-mutated modules in a bimodular setup *in vitro*, we yielded a large body of data giving insight into mechanistic details of the KS involved reactions. The amount of data was further increased by analyzing 8 selected substrate-accepting modules in combination with 3 novel substrate-donating modules. This allowed us to additionally analyze the influence of substrate-identity as well as domain-domain interactions during chain transfer (Figure 5A). A few trends emerged from this data. For example, (i) the chimeric bimodular PKSs using M3-TE generally show higher turnover rates when the native ACP2:KS3 interface is retained (except for Mut15), and (ii) the KS at the non-cognate module-module boundary in the chimeric bimodular systems represents a bottleneck that can be improved by mutating the binding site of the KS. Further, this data confirm previous reports of wild-type M3 preferring NDK over EDK,^16^ but this preference can be shifted by generating multipoint mutations. The usage of ACP2-NDK, reconstituting the native ACP2:KS3 interface, improved the efficiency of the system significantly (Figure 4B).

**Figure 5.**
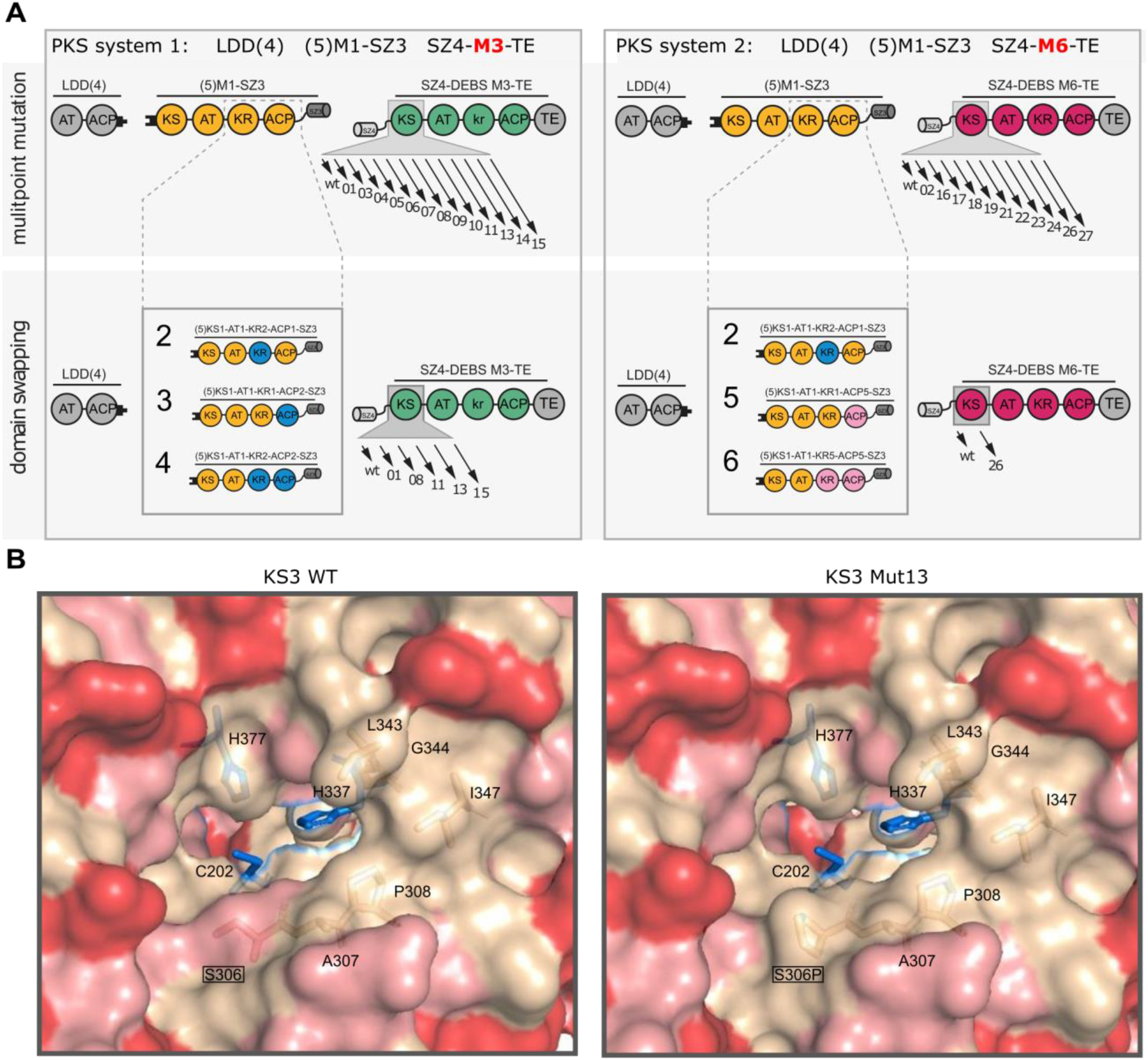
Systematic approach to analyze the parameters involved in the KS-mediated reactions and structural organization of the hydrophobic patch. (A) Systematic *in vitro* analysis of bimodular chimeric PKS systems comprised of LLD(4), (5)M1-SZ3 and SZ4-M3-TE (left) or SZ4-M6-TE (right). The interrogated KS-mutated modules are indicated with their numbering (e.g. 01 = Mut01). The novel substrate-donating modules are depicted. (B) Structural organization of the hydrophobic patch (A307, P308, L343, G344 and I347) for wildtype KS3 (left) and Mut13 (right) is depicted. The hydrophobic patch is suggested to serve as docking site for the intramodular ACP during chain translocation.^18^ The catalytic triad (C202, H337 and H377) is indicated in blue sticks. Charged residues are colored in red, polar residues in salmon, and hydrophobic residues in wheat.

While we observe complex and partly non-intuitive interplay of the factors constraining turnover rates in our data, we are able to give a mechanistic explanation for our observations in two cases, both of which can support PKS engineering approaches:

1. By taking a closer look at the different M3 construct designs, one specific mutation turns out to be putatively critical for increasing turnover rates. Intriguingly, from the 12 chosen designs, Mut08, Mut11, and Mut13, all among the highest efficient bimodular systems (Figure 3A), include the S306P mutation (Figure 2C, Figure S3). S306 resides in a loop that protrudes into the KS binding funnel. This loop is conformationally flexible, judged from crystallographic B-factors, and the S306 finds phi/psi angles that are not allowed for proline (see PDB entry code 2QO3)^19^. We assume that the S306P mutation changes the structural and conformational properties of the loop that can explain functional characteristics of the bimodular systems. Residue S306 is also located between the docking sites of the upstream and the intramodular ACP, that have recently been suggested for the homologous PIKS KS5 domain (Figures 5B&S12).^47^ While an *in vitro* mutagenesis study with DEBS KS1 could not support an impact of the S306 equivalent residue on the translocation reaction, it was suggested that the primary role of this residue is to stabilize the methylmalonyl-ACP bound to the KS binding site during elongation.^48^ In support of S306P affecting the elongation reaction, docking studies revealed a conserved hydrophobic patch near the entrance to the KS active site (formed by the residues A307, P308, L343, G344 and I347; KS3 numbering) that could interact with the intramodular ACP (Figures 5B&S12).^18^ Thus, the enlargement of the hydrophobic patch by the S306P mutation could lead to improved KS:ACP interactions during chain elongation. In conclusion, S306P may improve efficiencies of our chimeric bimodular systems by (i) releasing the specificity of the KS binding site by changed loop properties induced by the conformational rigidity of proline, and/or (ii) modulating the ACP:KS interfaces during chain translocation and elongation. The broad impact of S306P is in line with our data revealing improved turnover rates compared to the wildtype acceptor-module regardless of the employed substrate-donating module. We note that Mut15, that does not contain the proline mutation, behaved differently in this context. In addition, we would like to mention that turnover rates determined in this study do not *per se* differentiate between a kinetic barrier induced by the KS domain occurring during the translocation or the elongation step. Likely, KS mutations buried in the interior already affect the translocation step during the formation of the substrate-enzyme complex, and we follow that most of the KS mutations set in this study will address the translocation step. The residue S306 is located on the surface of KS3 and it is plausible that, as discussed above, this specific mutation could impact both the translocation and the elongation step.
2. We are also able to propose a molecular basis for the strikingly high efficiency of Mut26. M6-TE has been recognized before as suited substrate-accepting module for DEBS M1 in engineering approaches.^2,4,22,44,49^ Previously, the high sequence similarity between KS2 and KS6 has served as an explanation for wild-type M6 being almost as good as an acceptor as M2 with M1 as the substrate-donating module.^22^ As such, we expected that multipoint mutagenesis of KS6 would rather result in mutants with decreased activity with M1. Indeed, most of the screened mutants exhibited lower turnover numbers than wild-type M6, except for the multipoint mutant Mut26 showing intriguingly high turnover rates (Figure 3B). From structural inspection of the binding site of Mut26 we posit that the F235M mutation has likely the strongest impact. By replacing F235 with the smaller methionine, this mutation enlarges the binding pocket. In addition, as a more general consideration, the F235M mutation also increases the similarity between KS2 and KS6 even further (Figure S13). Additionally, residue V207 may be of relevance, because it resides in a hydrophobic pocket that is predicted to interact with the untethered end of the polyketide chain in DEBS KS3 and DEBS KS5.^19,42^ Generation of the V207L mutation in Mut26 may result in size reduction of this pocket, thereby leading to improved hydrophobic interactions with the smaller non-native substrate (diketide instead of pentaketide) donated by the upstream module in the chimeric assembly line. This data supports the assumption that the inherent KS substrate specificity represents a bottleneck in chimeric assembly lines and emphasizes the utility of KS mutagenesis to improve efficiencies.

## Conclusion

Beyond the boundaries of the DEBS PKSs that has served as a model system in this study, our data suggests limitations in engineering modular chimeric PKSs in general. As a yet unmatched systematic *in vitro* analysis of parameters involved in the KS-mediated uptake and processing of substrates (as part of the translocation and elongation reactions), our study can highlight why conventional engineering strategies have failed in the past.^2–4^ Further, our study indicates that also modern engineering approaches using the “updated module boundaries”^50^ will be limited in success as the revealed substrate specificities of the KS domains constitute crucial bottlenecks within chimeric assembly lines. In most cases, turnover efficiency can neither be deduced from the identity of the ACP donor, nor from the incoming substrate, as the highest turnover rates cannot be correlated with either of these parameters. Even worse from a perspective of simple mix-and-match engineering approaches, it appears that a cumulated effect and a non-intuitive interplay of the identity of both ACP and substrate influence the turnover of many downstream modules.

Engineering strategies are likely to be more successful when employing domains in chimeric PKS design that natively process chemically similar substrates^51,52^ and preserving cognate or installing cognate-like domain:domain interfaces,^36,53–55^ a task that can be achieved with the support from evolutionary analysis^34^ and computational PKS design.

## METHODS

### Reagents

CloneAmp HiFi PCR Premix was from Takara. Restriction enzymes were from New England Biolabs. All primers were synthesized by Sigma Aldrich. For DNA purification, the GeneJET Plasmid Miniprep Kit and the GeneJET Gel Extraction Kit from Thermo Scientific were used. Stellar Competent Cells were from Takara and One Shot^®^ BL21 (DE3) Cells were from Invitrogen. All chemicals for buffer preparations were from SigmaAldrich. Isopropyl-β-D-1-thiogalactopyranoside (IPTG), kanamycin sulfate, and carbenicillin were from Sigma Aldrich/Carl Roth. LB broth (Lennox) and 2xYT media for cell cultures were from Carl Roth, Ni-NTA affinity resin was from Clontech. For anion exchange chromatography the HiTrapQ column was from GE Healthcare. For protein concentration Amicon Ultra centrifugal filters were from Millipore. Coenzyme A (CoA), reduced β-nicotinamide adenine dinucleotide 2’-phosphate (NADPH), sodium propionate, methylmalonic acid, and magenesium chloride hexahydrate were from Carl Roth. Adenosine-5’-triphosphate (ATP) was from SigmaAldrich. Reducing agent tris(2-carboxyethyl)-phosphine (TCEP) was from Thermo Scientific. UV-Star half area microtiter plates were from Greiner.

### Plasmids

Plasmids harboring genes encoding individual PKS modules were either generated in this study via In-Fusion Cloning (Takara), site directed mutagenesis, or reported previously. Tables S6-S9 specify the cloning strategy, primer sequences, and the resulting plasmids. Plasmids sequences were verified by DNA sequencing (SeqLab). Protein sequences of substrate-donating modules newly generated in this study are presented in Table S10.

### Bacterial Cell Culture and Protein Purification

All PKS proteins were expressed and purified using similar protocols. For *holo*-proteins (where the ACP domain is post-translationally modified with a phosphopantetheine arm) *E. co/i* BL21 cells were co-transformed with a plasmid encoding for the phosphopantetheine transferase Sfp from *B. subtilis* (pAR3 5 7^56^). All proteins contained a C-terminal His_+_-tag for purification. Cultures were grown on a 1-2 L scale in 2xYT media at 37 °C to an OD_600_ of 0.3, whereupon the temperature was adjusted to 18 °C. At OD_600_ of 0.6, protein production was induced with 0.1 mM IPTG, and the cells were grown for another 18 h. Cells were harvested by centrifugation at 5000 g for 15 min and lysed by French Press (lysis buffer: 50 mM sodium phosphate, 10 mM imidazole, 450 mM NaCl, 10% glycerol, pH 7.6). The cell debris was removed by centrifugation at 50,000 g for 50 min. The supernatant was further purified using affinity chromatography (Ni-NTA agarose resin, 5 mL column volumes). After applying the supernatant to the column, a first wash step was performed with the above lysis buffer (10 column volumes), followed by a second wash step using 10 column volumes of wash buffer (50 mM phosphate, 25 mM imidazole, 300 mM NaCl, 10 % glycerol, pH 7.6). Proteins were eluted with 6 column volumes elution buffer (50 mM phosphate, 500 mM, 10% glycerol, pH 7.6). The eluate was further purified by anion exchange chromatography using a HitrapQ column on an ÄKTA FPLC system. Buffer A consisted of 50 mM phosphate, 10% glycerol, pH 7.6, whereas buffer B contained 50 mM phosphate, 500 mM NaCl, 10% glycerol, pH 7.6. The yields of all protein per liter of culture are noted in Table S2. Enzymes MatB, SCME, and PrpE were purified as decribed.^39,57^ Protein concentrations were determined with the BCA Protein Assay Kit (Thermo Scientific). Samples were stored as aliquots at −80 °C until further use.

### Size Exclusion Chromatography

To determine the purity of the proteins after ion exchange chromatography, samples of each protein were analyzed by size exclusion chromatography on an ÄKTA FPLC system using a Superose 6 Increase 10/300 GL column (buffer: 50 mM phosphate, 500 mM NaCl, 10% glycerol, pH 7.6). For protein purity by SDS-PAGE and SEC profiles refer to Figures S6, S7&S10.

### Thermal Shift Assay

The melting temperatures of all proteins in a buffer consisting of 50 mM phosphate, 500 mM NaCl, 10% glycerol, pH 7.6 were measured according to published procedures using a protein concentration of 1.2 mg/mL.^58^

### PKS Enzymatic Assays

PKS enzymatic assays were performed according to published procedures.^4^ PKS enzymes were used at 4 μM final concentration, for *in situ* substrate generation enzymes MatB, PrpE, and SCME were used at final concentrations of 4 μM, 2 μM, and 8 μM, respectively.

### Liquid Chromatography-Mass Spectrometry Analysis of Polyketides

Dried samples were reconstituted in 100 μL methanol, separated on a Acquity UPLC BEH C18 1.7 μm RP 2.1 x 50 mm column (Waters) with an Acquity UPLC BEH C18 1.7 μm RP 2.1 x 5 mm pre-column (Waters), connected to an Ultimate 3000 RSLC or an Ultimate 3000 LC (Dionex) HPLC over a 16 min linear gradient of acetonitrile from 5% to 95% in water, and subsequently injected into an Impact II qTof (Bruker) or an AmaZonX (Bruker) equipped with an ESI Source set to positive ionization mode. Reduced and unreduced triketide products were located by searching for the theoretical *m/z* for the [M+H]^+^-ion. Unreduced triketides: [M+H]^+^ = 171.098, reduced triketides [M+H]^+^ =173.118.

### Bioinformatical Analysis

Sequence alignments were generated with ClustalWS as implemented in *Jalview* 2.10.5.^59^ Homology models were generated with SWISS-MODEL^19^ and visualized PyMol. The FuncLib webserver was used to calculate diverse multipoint mutations within the KS binding site.^35^ The calculation for KS3 was based on the X-ray structure of the KS3-AT3 didomain.^42^ The calculation for KS6 was based on the homology model generated with SWISS-MODEL based on the X-ray structure of KS5-AT5.^19^ In the first step, for the calculation of the sequence space, chains A+B were included for KS3. To decrease the calculation effort only chain B was included for KS6. Seven residues were chosen for mutagenesis of KS3 and nine positions for KS6. In addition, residues of the catalytic triad (KS3: Cys202, H337 and H377; KS6: Cys172, H307 and H347) were selected not to be altered during simulations. The nomenclature of residues is derived from structural models. C202_KS3_ corresponds to UniProt EryA2 (Q03132) position 202 and C172_KS6_ to UniProt EryA3 (Q03133) position 1661. For generation of the multiple sequence alignment the default parameters were used during all calculations (Min ID 0.3, Max targets 3000, Coverage 0.6, and E value 0.0001). In the second calculation step, the parameters were chosen to yield between 500-500,000 designs. For KS3 152,826 and for KS6 285,822 designs harboring 3-5 mutations were selected for Rosetta calculation.

## Acknowledgments

We thank Helge Bode for use of their LC-MS instruments. We further want to thank Aleksandra Nivina and Chaitan Khosla for helpful discussions. In using the FuncLib method, we received fantastic support from Sarel Fleishman, Rosalie Lipsh and Jaime Prilusky.

## Authors contribution

The manuscript was written through contributions of all authors. M.K. conceived and supervised the project. M.K. cloned and purified alternative substrate-donating modules and analyzed the activity. L.B. designed the mutants, cloned and purified all mutant constructs and analyzed the activity, thermal stability, and oligomerization. M.G. designed the research and analyzed the data.

## Supporting Information Available

Sequence alignments, structural models, SEC data, SDS-PAGE data, LC-MS data, FuncLib parameters, list of plasmids, construct design, protein sequences, and expression yields. This material is available free of charge via the internet at http://pubs.acs.org.

