## Supplemental Information for "The Ketosynthase Domain Constrains the Design of Polyketide Synthases"

**Content:**

**Supplementary Figures:**

Figure S1. Sequence alignment of KS domains to identify residues for site-directed mutagenesis of DEBS KS3.
Figure S2. Sequence alignment of KS domains to identify residues for site-directed mutagenesis of DEBS KS6.
Figure S3. Structural alignment of wild-type and mutant KS3 binding sites.
Figure S4. Structural alignment of wild-type and mutant KS6 binding sites.
Figure S5. Purity of proteins used in this study.
Figure S6. Analysis of proteins by SEC – M3 substrate-accepting modules.
Figure S7. Analysis of proteins by SEC – M6 substrate-accepting modules.
Figure S8. LC-MS analysis of unreduced triketide lactone 2 produced by chimeric bimodular PKSs using (5)M1-SZ3 and SZ4-M3-TE.
Figure S9. LC-MS analysis of reduced triketide lactone 3 and unreduced triketide lactone 5 produced by chimeric bimodular PKSs using (5)M1-SZ3 and SZ4-M6-TE.
Figure S10. Analysis of substrate-donating modules by SEC.
Figure S11. LC-MS analysis of unreduced triketide lactone 2 (in case of Donor 3) and unreduced triketide lactone 4 (in case of Donor 2 & 4) produced by chimeric bimodular PKSs using SZ4-M3-TE and different substrate-donating modules.
Figure S12. Structural organization of ACP docking interfaces and hydrophobic patch in DEBS KS3 and KS3_Mut13.
Figure S13. Sequence and structural alignment of DEBS KS2 and KS6.

**Supplementary Tables:**

Table S1. Sequence space of chosen binding site residues of KS3 and KS6 calculated by FuncLib
Table S2. Yields of proteins used in this study
Table S3. Melting temperatures of wild-type and mutant modules
Table S4. Distribution of reduced and unreduced triketide lactone in bimodular chimeric PKSs using LDD(4), (5)M1-SZ3 and the listed substrate-accepting modules.
Table S5. Peak area of reduced and unreduced triketide lactone products in bimodular chimeric PKSs using M6-TE or Mut26 in the presence of different substrate-donating modules.
Table S6. Plasmids used in this study and their origin
Table S7. Cloning strategy of plasmids generated in this study
Table S8. Cloning strategy of for fragments used in In-Fusion cloning and primer sequences
Table S9. gBlocks used for In-Fusion cloning
Table S10. Amino acid sequences of newly generated substrate-donating modules

### Supplementary Figures

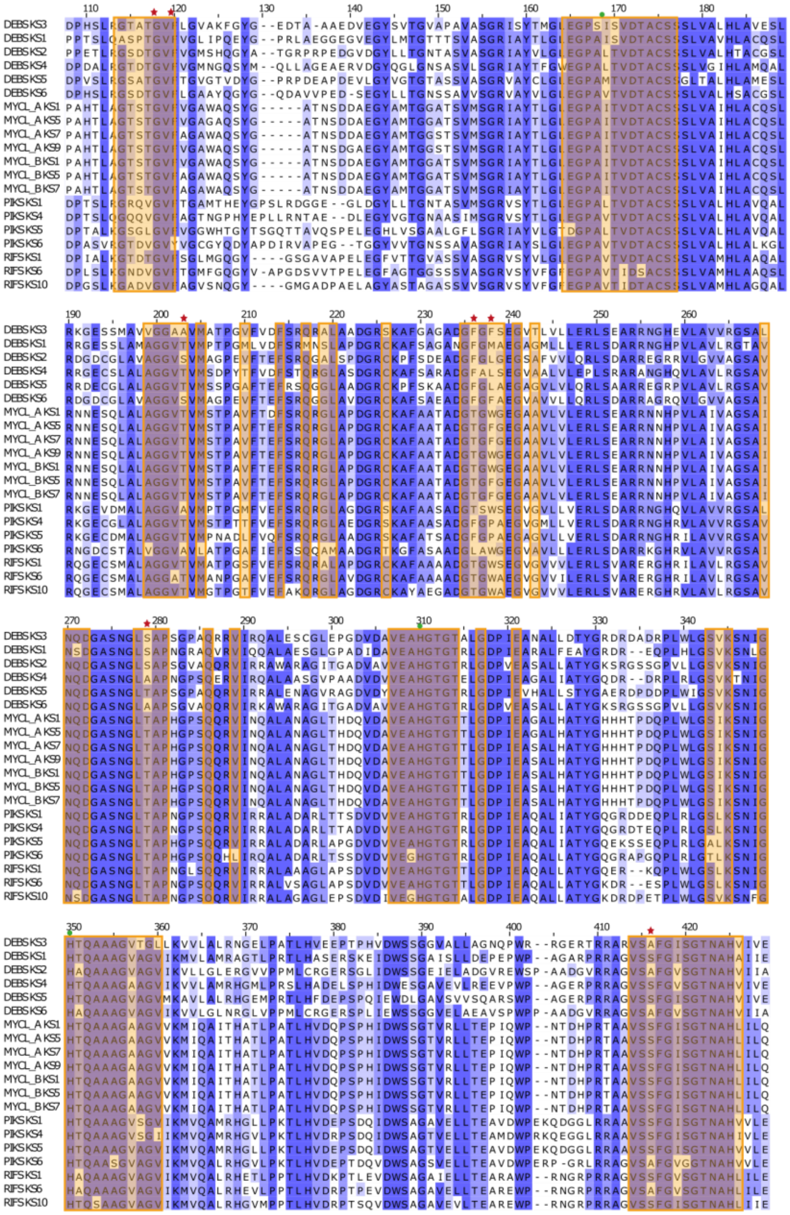

**Figure S1. Sequence alignment of KS domains to identify residues for site-directed mutagenesis of DEBS KS3.** Sequences were obtained from DEBS, mycolactone synthase (MYCL), PIKS, and RIFS. Residues within 12 Å of the active cysteine of DEBS KS3 (C175; Uniprot EryA2 Q03132 position 202) are highlighted in orange. Green circles- catalytic triad, red stars- residues selected for multipoint mutagenesis (A124, F126, A203, F236, F238, S279, and A416). Nomenclature of selected residues according to PDB 2OQ3: A154, F156, A230, F263, F265, S306, A441.

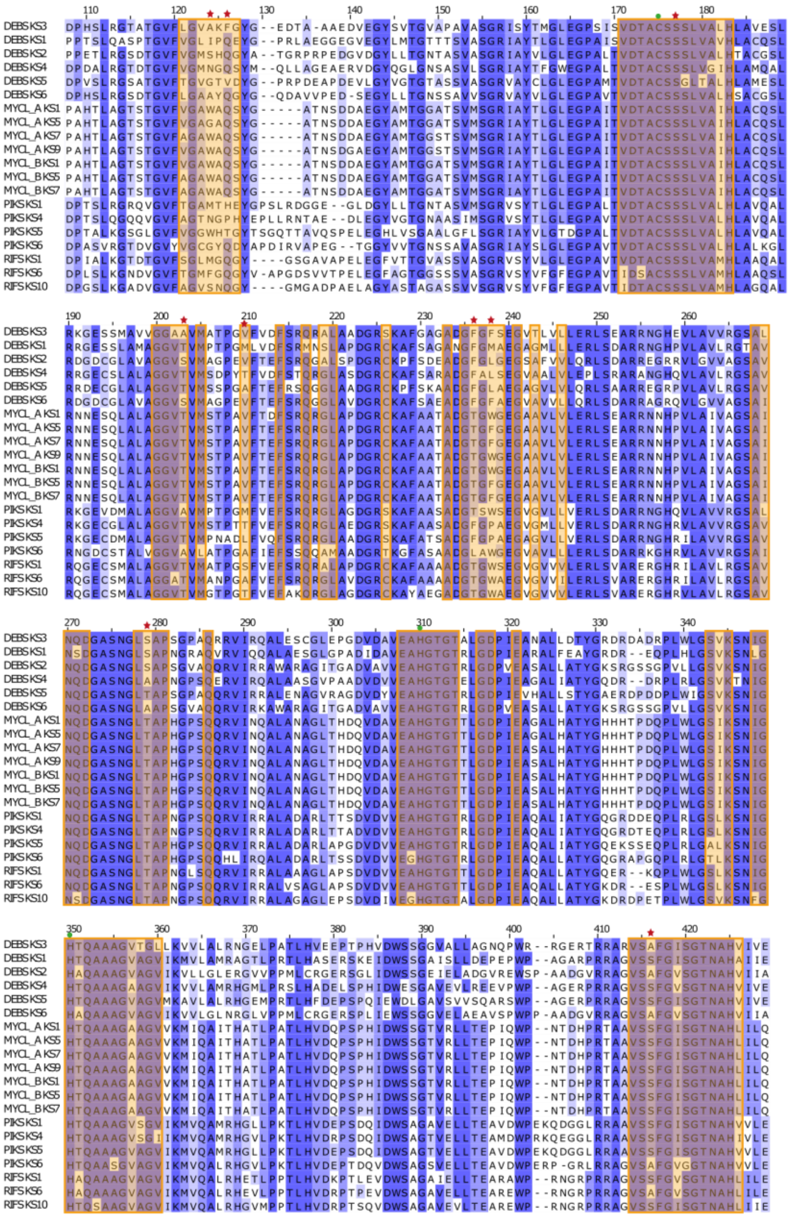

**Figure S2. Sequence alignment of KS domains to identify residues for site-directed mutagenesis of DEBS KS6.** Sequences were obtained from DEBS, mycolactone synthase (MYCL), PIKS, and RIFS. Residues within 12 Å of the active cysteine of DEBS KS6 (C175; Uniprot EryA3 Q03133 position 1661) are highlighted in orange. Green circles- catalytic triad, red stars- residues selected for multipoint mutagenesis (A124, Q126, S177, S203, V210, F236, F238, A279, A416). Nomenclature of selected residues according to homology model: A124, Q126, S174, S200, V207, F233, F235, A276, A412. Note that S177 (S174 in homology model) was accidently included in the diversified sequence space, although it is a conserved residue. According to the phylogenetic analysis implemented in FuncLib both Ala, and Gly can also be tolerated at this position (see Table S1).

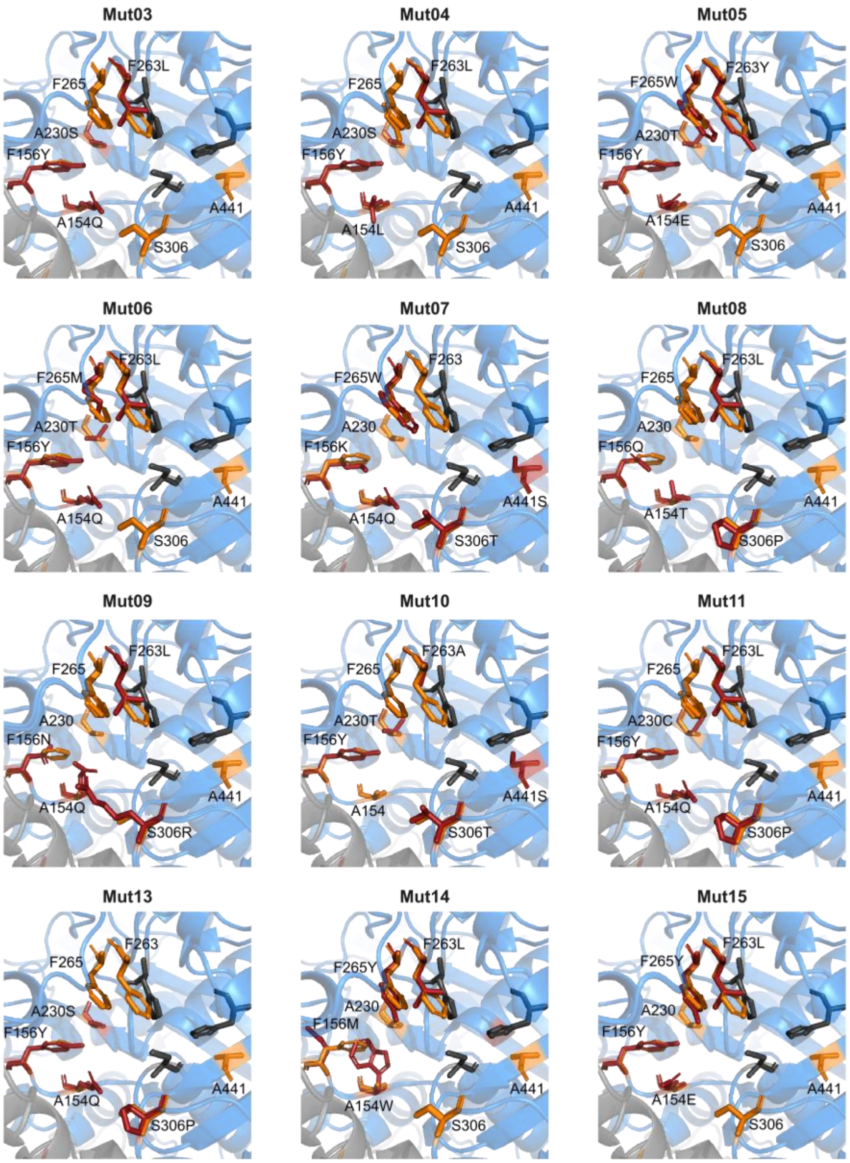


**Figure S3. Structural alignment of wild-type and mutant KS3 binding sites.** Alignment of wild-type KS3 (PDB 2QO3) and predicted structures of mutant KS3 proteins. Catalytic triad in gray, residues selected for mutagenesis in orange, and residues mutant in the respective design in red.

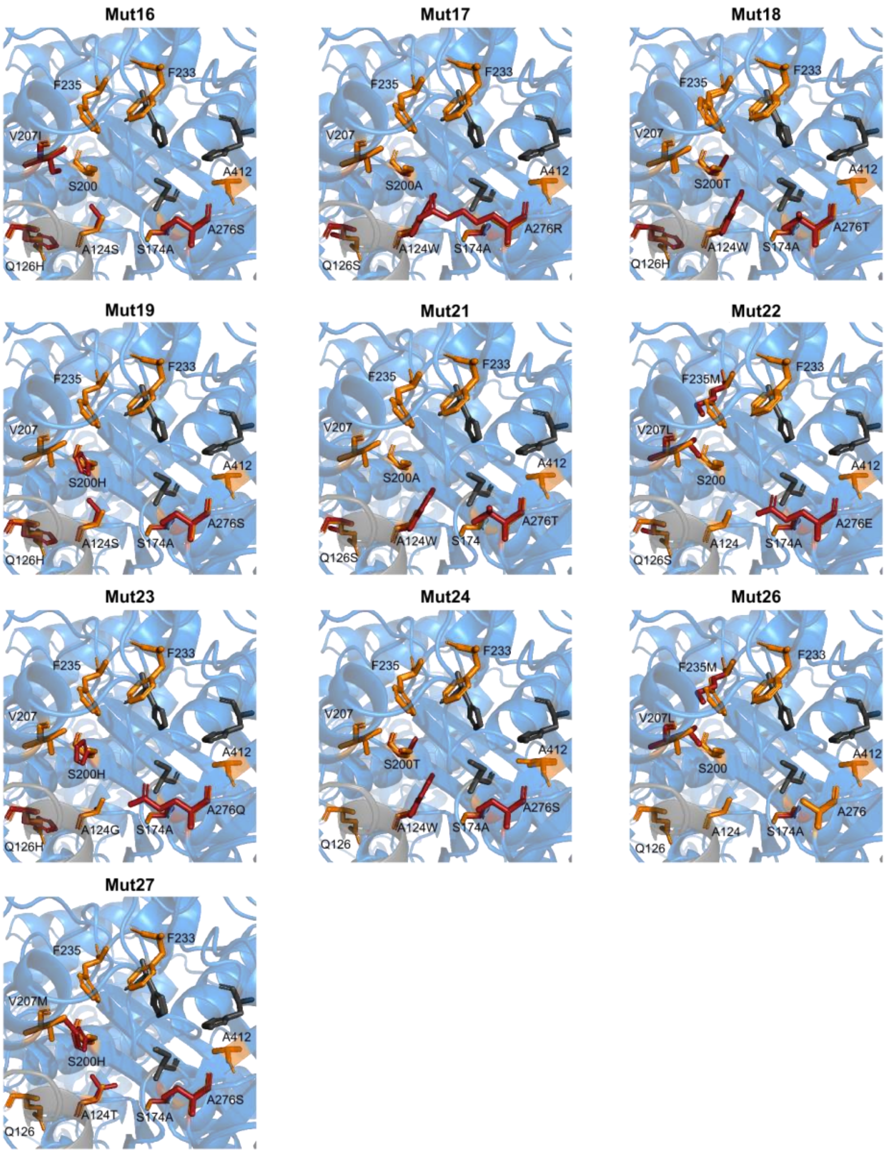


**Figure S4. Structural alignment of wild-type and mutant KS6 binding sites.** Alignment of wild-type KS6 (homology model) and predicted structures of mutant KS6 proteins. Catalytic triad in gray, residues selected for mutagenesis in orange, and residues mutant in the respective design in red.

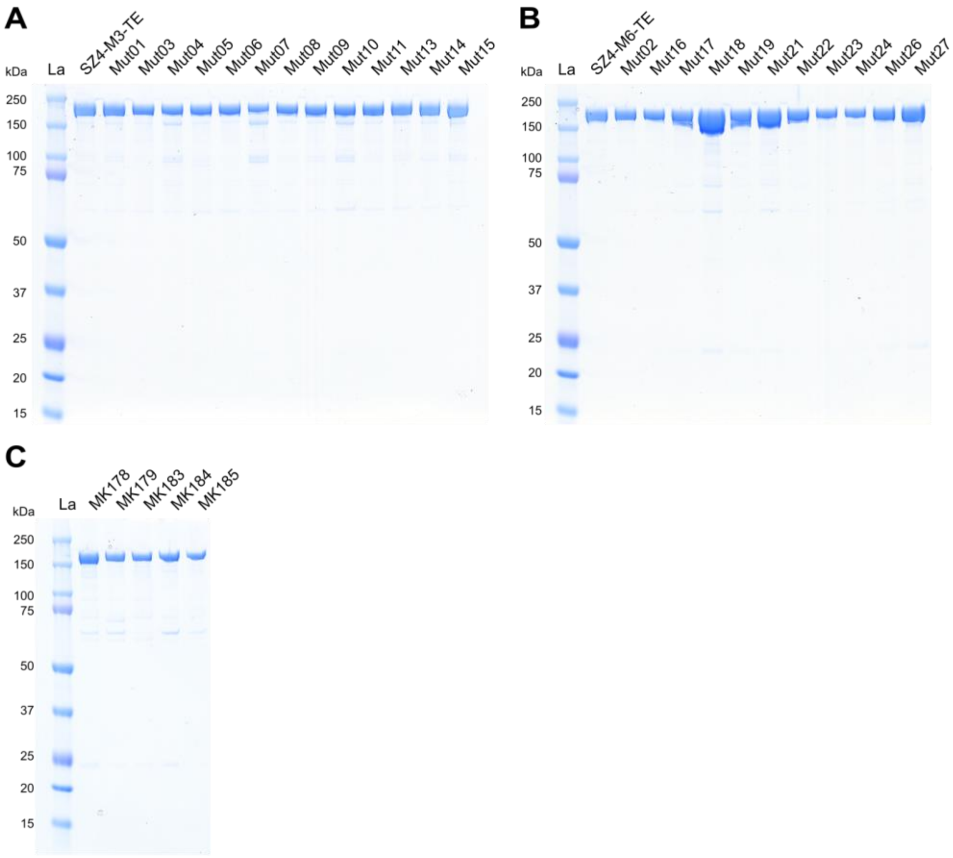


**Figure S5. Purity of proteins used in this study.** (A) SZ4-M3-TE and its mutants (MW 189 kDa), (B) SZ4-M6-TE and its mutants (MW 184 kDa), and (C) substrate-donating modules MK178 (Donor 4, 161 kDa), MK179 (Donor 6, 163 kDa), MK183 (Donor 2, 161 kDa), MK184 (Donor 3, 164 kDa), MK185 (Donor 5, 164 kDa).

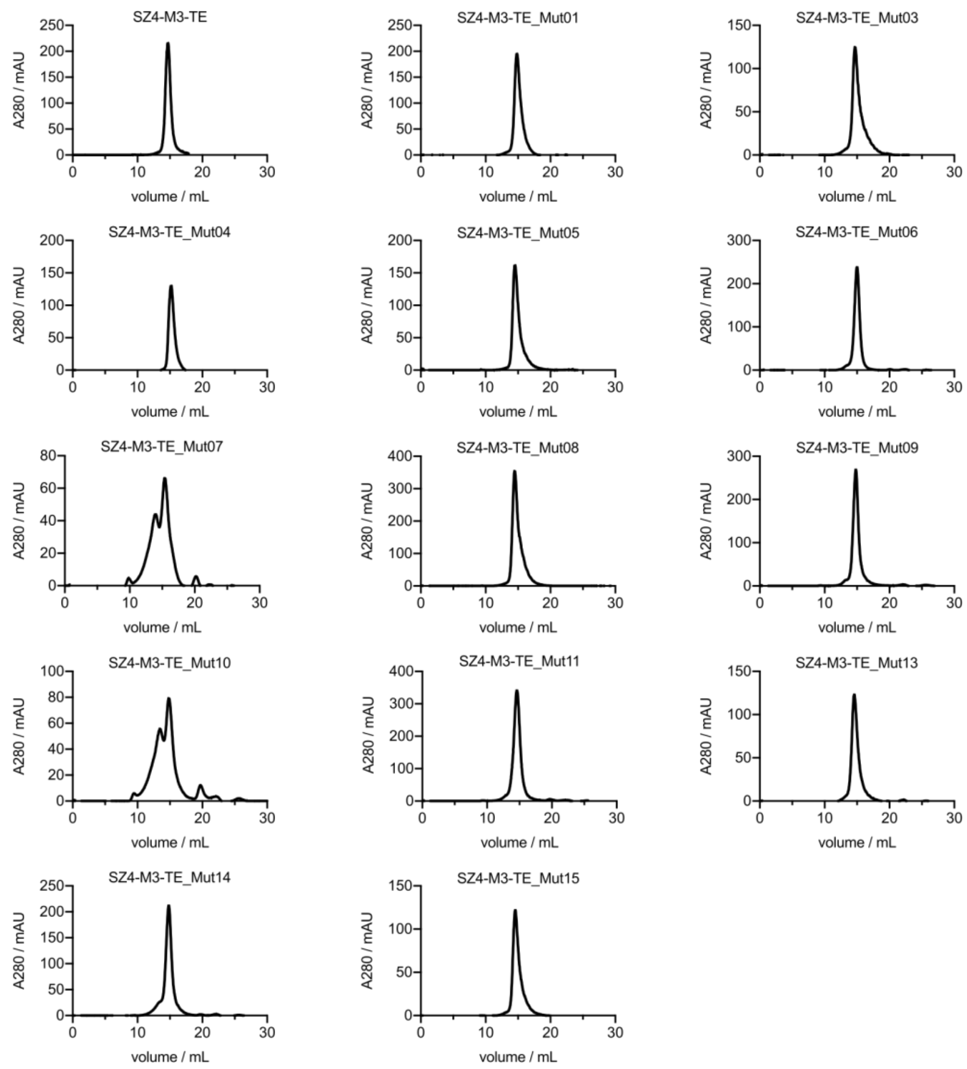


**Figure S6. Analysis of proteins by SEC – M3 substrate-accepting modules.** Wild-type M3-TE and its mutants analyzed by SEC. All proteins eluted in a predominately single peak from SEC.

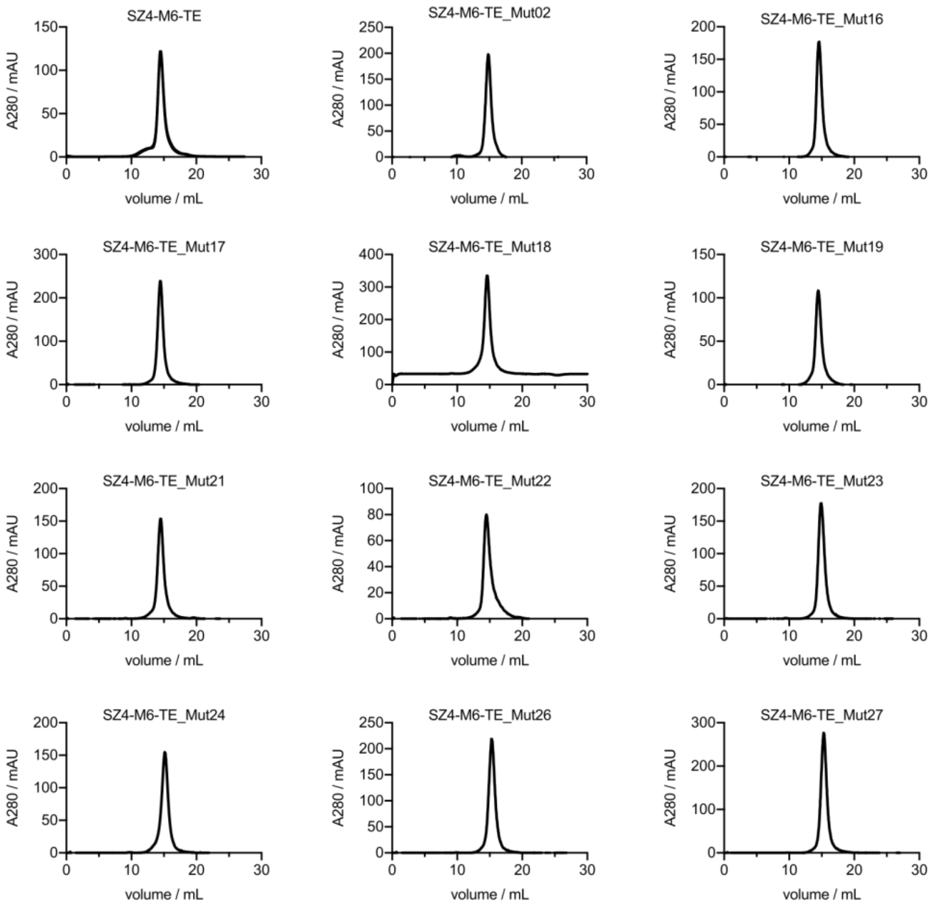


**Figure S7. Analysis of proteins by SEC – M6 substrate-accepting modules.** Wild-type M6-TE and its mutants analyzed by SEC. All proteins eluted in a predominately single peak from SEC.

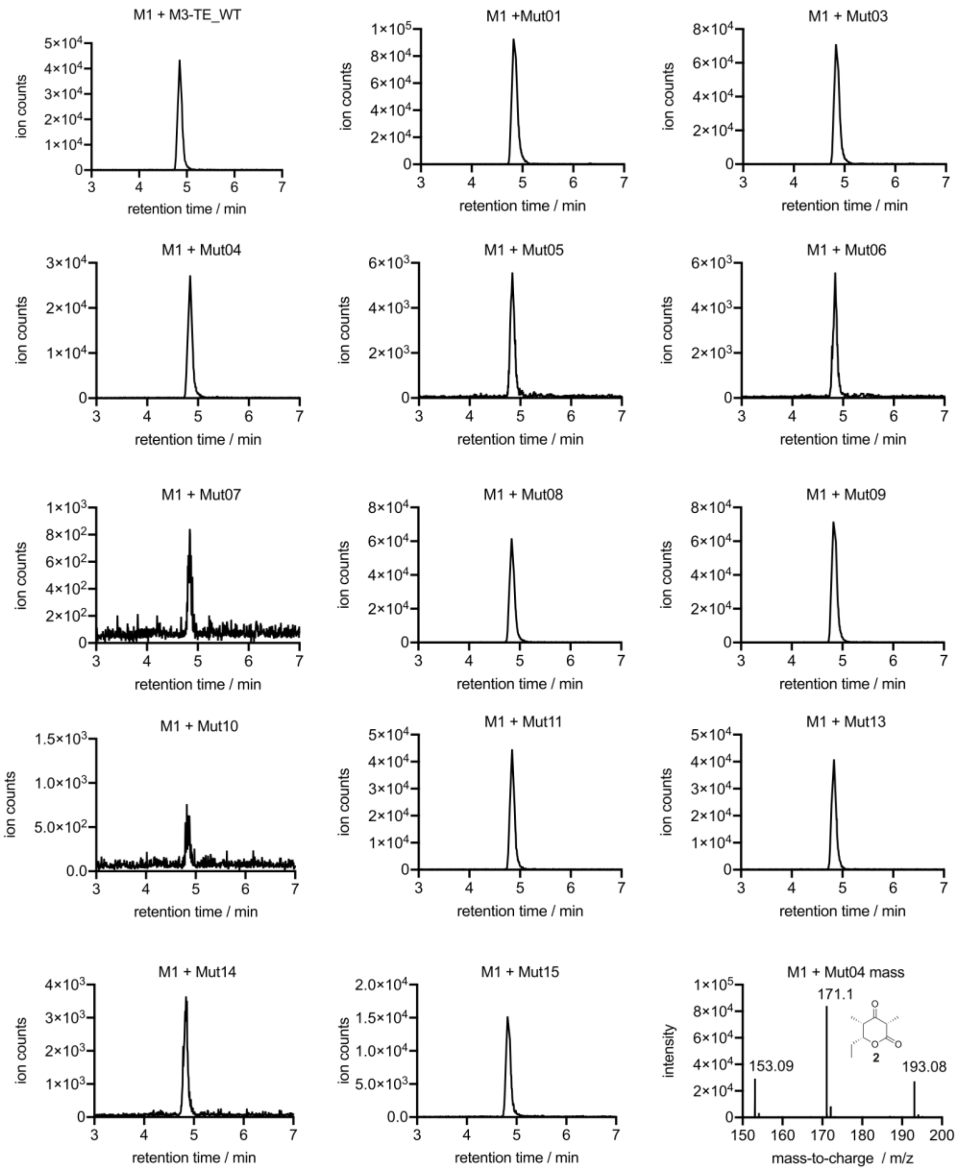


**Figure S8. LC-MS analysis of unreduced TKL 2 produced by chimeric bimodular PKSs using (5)M1-SZ3 and SZ4-M3-TE.** Unreduced TKL **2** ($C_9H_{14}O_3$, calculated MW = 170.22 g/mol) was detected in all reaction mixtures after overnight incubation. The extracted ion chromatograms were obtained by extraction of the $[M+H]^+$ species and one chromatogram per compound is shown as an example. Labeled peaks from left to right correspond to $[M+H-H2O]^+$, $[M+H]^+$, and $[M+Na]^+$ ions. TKL **2** eluted at 4.7 min.

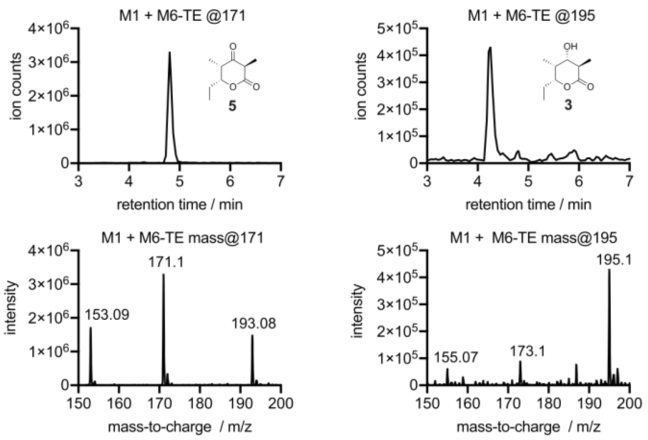


**Figure S9. LC-MS analysis of reduced TKL 3 and unreduced TKL 5 produced by chimeric bimodular PKSs using (5)M1-SZ3 and SZ4-M6-TE.** Reduced TKL **3** ($C_9H_{16}O_3$, calculated MW = 172.22 g/mol) and unreduced TKL **5** ($C_9H_{14}O_3$, calculated MW = 170.22 g/mol) were simultaneously detected in reaction mixtures shown in Figure 3B. The extracted ion chromatograms were obtained by extraction of the $[M+H]^+$ species (TKL **5**) or $[M+Na]^+$ species (TKL **3**) and one chromatogram per compound is shown as an example. Labeled peaks from left to right correspond to $[M+H-H2O]^+$, $[M+H]^+$, and $[M+Na]^+$ ions. TKL **5** eluted at 4.7 min and TKL **3** at 4.2 min.

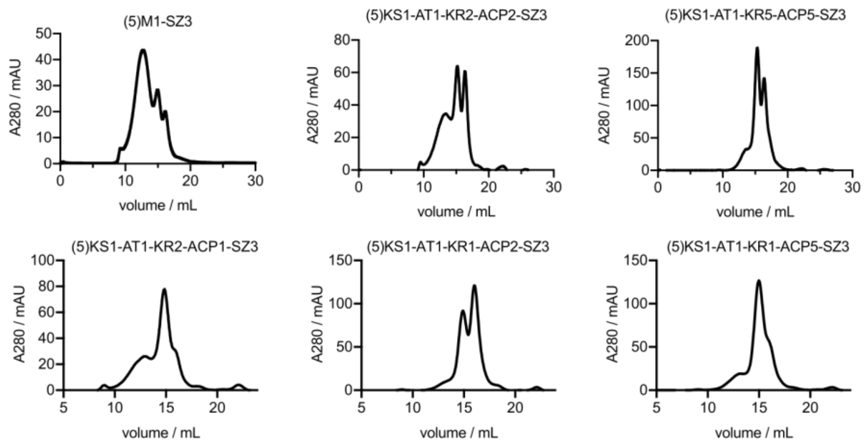


**Figure S10. Analysis of substrate-donating modules by SEC.** All proteins eluted in multiple oligomeric species from SEC.

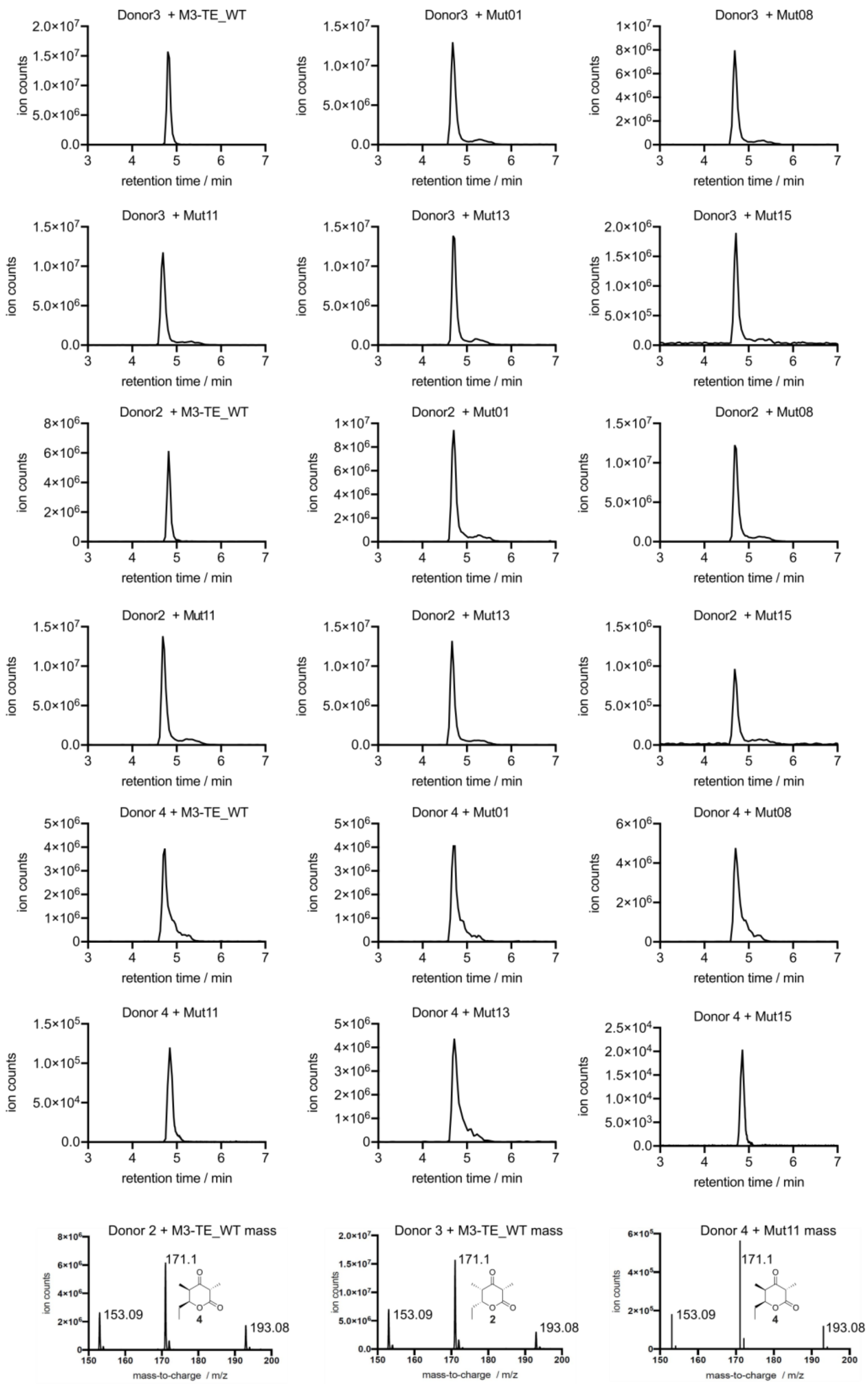

**Figure S11. LC-MS analysis of unreduced TKL 2 (in case of Donor 3) and unreduced TKL 4 (in case of Donor 2 & 4) produced by chimeric bimodular PKSs using SZ4-M3-TE and different substrate-donating modules.** Either (5)KS1-AT1-KR1-ACP2-SZ3 (Donor 3), (5)KS1-AT1-KR2-ACP1-SZ3 (Donor 2), or (5)KS1-AT1-KR2-ACP2-SZ3 (Donor 4) was used as the substrate-donating module**.** Unreduced TKL **2** or the diastereomeric form unreduced TKL **4** ($C_9H_{14}O_3$, calculated MW = 170.22 g/mol) was detected in all reaction mixtures after overnight incubation. The extracted ion chromatograms were obtained by extraction of the $[M+H]^+$ species and one chromatogram per substrate-donating module is shown as an example. Labeled peaks from left to right correspond to $[M+H-H2O]^+$, $[M+H]^+$ and $[M+Na]^+$ ions.

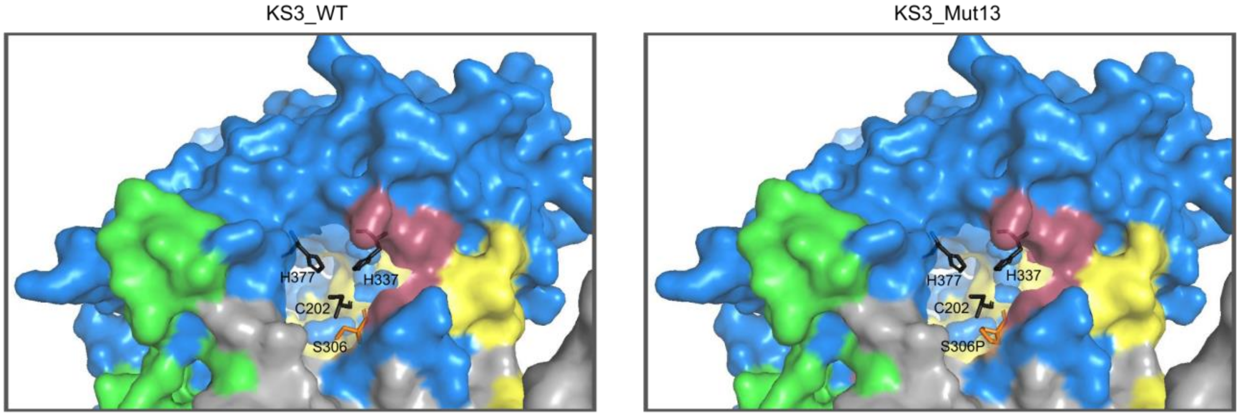


**Figure S12. Structural organization of ACP docking interfaces and hydrophobic patch in DEBS KS3 and KS3_Mut13.** The catalytic triad is depicted as black sticks. Position S306 which enlarged the conserved hydrophobic patch when mutated to S306P is depicted as orange stick. Docking interfaces of the upstream and intramodular ACP[1] are shown as yellow and green surfaces, respectively. The hydrophobic patch[2] is indicated as purple surface. Chain A and chain B of the KS3-AT3 didomain are colored in blue and grey, respectively.

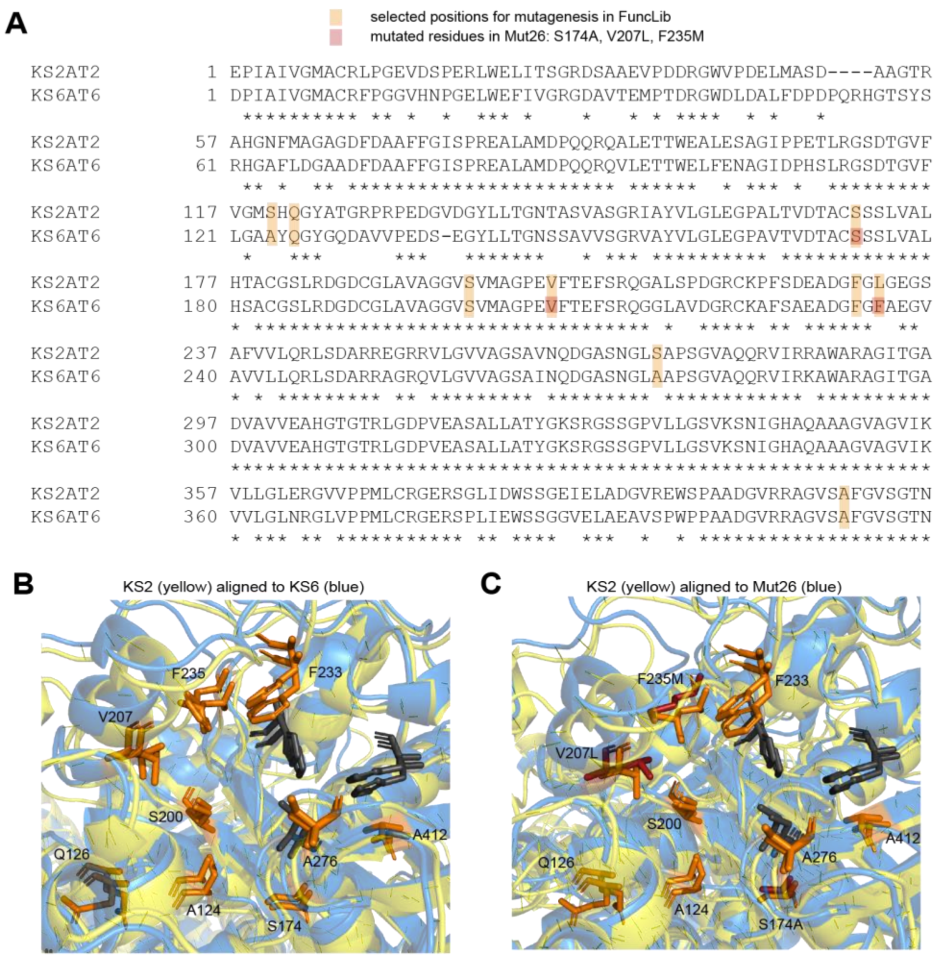


**Figure S13. Sequence and structural alignment of DEBS KS2 and KS6.** (A) Sequence alignment of KS2 and KS6. Residues selected for mutagenesis are highlighted in orange and those mutated in Mut26 in red. Structural alignment of KS2 to KS6 (B) and of KS2 to Mut26 (C). Catalytic triad in gray, residues selected for mutagenesis in orange, and positions mutated in Mut26 in red.

### Supplementary Tables

**Table S1. Sequence space of chosen binding site residues of KS3 and KS6 calculated by FuncLib**

| KS Domain | Position | Theoretical sequence space |
|---|---|---|
| KS3 | A154 | ADEGIKLMNQSTVW |
| | F156 | FADEGHKMNQSTY |
| | A230 | ACST |
| | F263 | FALSY |
| | F265 | FMWY |
| | S306 | SAEGKMNPQRTV |
| | A441 | AS |
| KS6 | A124 | AGNQSTW |
| | Q126 | QAGHNST |
| | S174 | SAG |
| | S200 | SAHMNT |
| | V207 | VACHILMNQST |
| | F233 | FAS |
| | F235 | FM |
| | A276 | AEGKNQRST |
| | A412 | ADEGS |

**Table S2. Yields of proteins used in this study.** Typical yields are presented.

| Construct | Protein | Yield/ mg/L of *E. coli* culture |
|---|---|---|
| MK149 | SZ4-M3-TE | 5.9 |
| LB001 | SZ4-M3-TE_Mut01 | 7.9 |
| LB003 | SZ4-M3-TE_Mut03 | 11.0 |
| LB004 | SZ4-M3-TE_Mut04 | 5.4 |
| LB005 | SZ4-M3-TE_Mut05 | 7.2 |
| LB006 | SZ4-M3-TE_Mut06 | 8.5 |
| LB007 | SZ4-M3-TE_Mut07 | 8.4 |
| LB008 | SZ4-M3-TE_Mut08 | 7.9 |
| LB009 | SZ4-M3-TE_Mut09 | 8.6 |
| LB010 | SZ4-M3-TE_Mut10 | 11.0 |
| LB011 | SZ4-M3-TE_Mut11 | 17.0 |
| LB013 | SZ4-M3-TE_Mut13 | 5.7 |
| LB014 | SZ4-M3-TE_Mut14 | 10.0 |
| LB015 | SZ4-M3-TE_Mut15 | 6.9 |
| MK147 | SZ4-M6-TE | 6.3 |
| LB002 | SZ4-M6-TE_Mut02 | 5.5 |
| LB016 | SZ4-M6-TE_Mut16 | 9.1 |
| LB017 | SZ4-M6-TE_Mut17 | 11.0 |
| LB018 | SZ4-M6-TE_Mut18 | 11.0 |
| LB019 | SZ4-M6-TE_Mut19 | 8.2 |
| LB021 | SZ4-M6-TE_Mut21 | 9.6 |
| LB022 | SZ4-M6-TE_Mut22 | 5.1 |
| LB023 | SZ4-M6-TE_Mut23 | 6.2 |
| LB024 | SZ4-M6-TE_Mut24 | 6.5 |
| LB026 | SZ4-M6-TE_Mut26 | 7.2 |
| LB027 | SZ4-M6-TE_Mut27 | 7.9 |
| BL12 | LDD(4) | 2.9 |
| MK150 | (5)M1-SZ3 | 4.7 |
| MK148 | SZ4-M2-TE | 2.5 |
| MK178 | (5)KS1-AT1-KR2-ACP2-SZ3 | 4.1 |
| MK179 | (5)KS1-AT1-KR5-ACP5-SZ3 | 5.4 |
| MK183 | (5)KS1-AT1-KR2-ACP1-SZ3 | 2.7 |
| MK184 | (5)KS1-AT1-KR1-ACP2-SZ3 | 3.6 |
| MK185 | (5)KS1-AT1-KR1-ACP5-SZ3 | 6.5 |

**Table S3. Melting temperatures of wild-type and mutant modules.** N/A.- Not applicable due to higher oligomerization of protein in the sample (see Figure S6).

| Construct | $T_m$ / °C |
|---|---|
| MK147 | 41.7±0.3 |
| MK149 | 41.5±0.0 |
| LB001 | 40.0±0.0 |
| LB002 | 41.5±0.0 |
| LB003 | 41.7±0.3 |
| LB004 | 40.7±0.3 |
| LB005 | 41.5±0.0 |
| LB006 | 42.5±0.0 |
| LB007 | N/A |
| LB008 | 41.2±0.3 |
| LB009 | 43.7±0.3 |
| LB010 | N/A |
| LB011 | 41.5±0.7 |
| LB013 | 41.2±0.3 |
| LB014 | 46.0±0.0 |
| LB015 | 41.7±0.3 |
| LB016 | 41.2±0.3 |
| LB017 | 36.7±1.0 |
| LB018 | 40.7±0.3 |
| LB019 | 42.7±0.3 |
| LB021 | 41.0±0.0 |
| LB022 | 40.5±0.0 |
| LB023 | 38.5±0.0 |
| LB024 | 40.0±0.7 |
| LB026 | 40.2±0.3 |
| LB027 | 42.0±0.7 |

**Table S4. Distribution of reduced and unreduced triketide lactone products in bimodular chimeric PKSs using LDD(4), (5)M1-SZ3 and the listed substrate-accepting modules.** Peak areas were calculated from LC-MS measurements after the reaction time of 10 min. Peaks were obtained by searching for the respective $[M+H]^+$-species. Reduced TKL **3** eluted at 4.2 min and unreduced TKL **5** at 4.7 min.

| acceptor module | red. TKL, peak area | unred. TKL, peak area | red. TKL: unred. TKL |
|---|---|---|---|
| M2-TE | $2.3x10^7$ | $7.1x10^6$ | 3.2 : 1 |
| M6-TE_WT | $5.9x10^6$ | $1.7x10^8$ | 1 : 28.8 |
| M6-TE_Mut02 | $4.2x10^6$ | $1.1x10^8$ | 1 : 26.1 |
| M6-TE_Mut16 | $6.3x10^7$ | $2.0x10^8$ | 1 : 3.1 |
| M6-TE_Mut17 | $2.2x10^6$ | $1.0x10^8$ | 1 : 45.4 |
| M6-TE_Mut18 | $4.2x10^6$ | $1.3x10^8$ | 1 : 30.9 |
| M6-TE_Mut19 | $1.8x10^6$ | $5.4x10^7$ | 1 : 30.0 |
| M6-TE_Mut21 | $4.5x10^6$ | $1.2x10^8$ | 1 : 26.6 |
| M6-TE_Mut22 | $2.1x10^6$ | $3.9x10^7$ | 1 : 18.5 |
| M6-TE_Mut23 | $2.4x10^6$ | $6.4x10^7$ | 1 : 26.6 |
| M6-TE_Mut24 | $6.9x10^6$ | $2.3x10^8$ | 1 : 33.3 |
| M6-TE_Mut26 | $1.3x10^7$ | $5.1x10^8$ | 1 : 39.2 |
| M6-TE_Mut27 | $5.1x10^5$ | $3.0x10^7$ | 1 : 58.8 |

**Table S5. Peak area of reduced and unreduced triketide lactone products in bimodular chimeric PKSs using SZ4-M6-TE or Mut26 in the presence of different substrate-donating modules.** The identity of the substrate on the upstream ACP is indicated. Peak areas were calculated from LC-MS measurements after a reaction time of 10 min. Peaks were obtained by searching for the respective $[M+H]^+$-species. Reduced TKL **3** & **6** eluted at 4.2 min and unreduced TKL **5** & **7** at 4.7 min. Note that the ratio does not reflect the absolute amount of TKL product in the sample as no TKL standard was available to capture different ionization behaviors of the two compounds.

| Acceptor module | Donor module (Abbreviation and full name) | Substrate bound to upstream ACP | red. TKL, peak area | unred. TKL, peak area |
|---|---|---|---|---|
| M6-TE | 1, M1 | NDK | $5.9x10^6$ | $1.7x10^8$ |
| | 5, KS1-AT1-KR1-ACP5 | NDK | none | $5.8x10^7$ |
| | 2, KS1-AT1-KR2-ACP1 | EDK | $1.1x10^7$ | $6.4x10^6$ |
| | 6, KS1-AT1-KR5-ACP5 | EDK | $3.6x10^6$ | $2.6x10^6$ |
| M6-TE_Mut26 | 1, M1 | NDK | $1.3x10^7$ | $5.1x10^8$ |
| | 5, KS1-AT1-KR1-ACP5 | NDK | $3.1x10^6$ | $9.3x10^7$ |
| | 2, KS1-AT1-KR2-ACP1 | EDK | $1.1x10^7$ | $1.5x10^6$ |
| | 6, KS1-AT1-KR5-ACP5 | EDK | $1.8x10^7$ | $2.1x10^6$ |

**Table S6. Plasmids used in this study and their origin**

| Plasmid | Origin |
|---|---|
| pBL12_LDD(4)_pET28_kan | Lowry *et al.*[3] |
| pMK147_SZ4-M6-TE-H6_pET22_carb | Klaus *et al.*[4] |
| pMK148_SZ4-M2-TE-H6_pET22_carb | Klaus *et al* [4] |
| pMK149_SZ4-M3-TE-H6_pET22_carb | Klaus *et al.*[4] |
| pMK150_(5)M1-SZ3-H6_pET22b_carb | Klaus *et al.*[4] |
| pMK178_(5)KS1-AT1-KR2-ACP2-SZ3-H6_pET22b_carb | this study |
| pMK179_(5)KS1-AT1-KR5-ACP5-SZ3-H6_pET22b_carb | this study |
| pMK183_(5)KS1-AT1-KR2-ACP1-SZ3-H6_pET22b_carb | this study |
| pMK184_(5)KS1-AT1-KR1-ACP2-SZ3-H6_pET22b_carb | this study |
| pMK185_(5)KS1-AT1-KR1-ACP5-SZ3-H6_pET22b_carb | this study |
| pLB001_SZ4-M3-TE-H6_A189W_pET22_carb | this study |
| pLB002_SZ4-M6-TE-H6_A189W_pET22_carb | this study |
| pLB003_SZ4-M3-TE-H6_A189Q_F191Y_A265S_F298L_pET22_carb | this study |
| pLB004_SZ4-M3-TE-H6_A189L_F191Y_A265S_F298L_F300Y_pET22_carb | this study |
| pLB005_SZ4-M3-TE-H6_A19E_F191Y_A165T_F298Y_F300W_pET22_carb | this study |
| pLB006_SZ4-M3-TE-H6_A189Q_F191Y_A265T_F298L_F300M_pET22_carb | this study |
| pLB007_SZ4-M3-TE-H6_A189Q_F191K_F298W_S341T_A476S_pET22_carb | this study |
| pLB008_SZ4-M3-TE-H6_A189T_F191Q_F298L_S306P_pET22_carb | this study |
| pLB009_SZ4-M3-TE-H6_A189Q_F191N_F298L_S341R_pET22_carb | this study |
| pLB010_SZ4-M3-TE-H6_F191Y_A265T_F298A_S341T_A476S_pET22_carb | this study |
| pLB011_SZ4-M3-TE-H6_A189Q_F191Y_A265C_F298L_S341P_pET22_carb | this study |
| pLB013_SZ4-M3-TE-H6_A189Q_F156Y_A265S_S341P_pET22_carb | this study |
| pLB014_SZ4-M3-TE-H6_A189W_F191M_F298L_F300Y_pET22_carb | this study |
| pLB015_SZ4-M3-TE-H6_A189E_F191Y_F298L_F300Y_pET22_carb | this study |
| pLB016_SZ4-M6-TE-H6_A189S_Q191H_S239A_V272I_A341S_pET22_carb | this study |
| pLB017_SZ4-M6-TE-H6_A189W_Q191S_S239A_S265A_A341R_pET22_carb | this study |
| pLB018_SZ4-M6-TE-H6_A189W_Q191H_S239A_S265T_A341T_pET22_carb | this study |
| pLB019_SZ4-M6-TE-H6_A189S_Q191H_S239A_S265H_A341S_pET22_carb | this study |
| pLB021_SZ4-M6-TE-H6_A189W_Q191S_S265A_A341T_pET22_carb | this study |
| pLB022_SZ4-M6-TE-H6_Q191S_S239A_V272L_F300M_A341E_pET22_carb | this study |
| pLB023_SZ4-M6-TE-H6_A189G_Q191H_S239A_S265H_A341Q_pET22_carb | this study |
| pLB024_SZ4-M6-TE-H6_A189W_S239A_S265T_A341S_A477S_pET22_carb | this study |
| pLB026_SZ4-M6-TE-H6_S239A_V272L_F300M_pET22_carb | this study |
| pLB027_SZ4-M6-TE-H6_A189T_S239A_S265H_V272M_A341S_pET22_carb | this study |

**Table S7. Cloning strategy of plasmids generated in this study.** Individual fragments were generated by overlap extension PCR, conventional PCR or ordered as gBlocks and assembled via In-Fusion cloning. .Single point mutations were introduced via QuickChange site-directed mutagenesis. See Table S8 for primer used to generate individual fragments and Table S9 for sequences of gBlocks.

| Plasmid | Cloning Method | Fragments | Primer Name | Primer Sequence 5'-3' | Template |
|---|---|---|---|---|---|
| pLB003 | In-Fusion | 1 + 7 | P-LB003 | CTCGCCGTAGCCGAACTTCcaCACTCCGAGGAAGACGCC | pMK149 |
| | | | P-LB050 | TCCGCTCGCCGCGCC | |
| | | 5 + 3 | P-LB051 | GCGCGAGGGCGGTGAG | pMK149 |
| | | | P-LB009 | CGGGGTCGCCATCACCGaGGCACCGCCGACGACC | |
| | | pLB003V | P-LB052 | CTCACCGCCCTCGCGC | pMK149 |
| | | | P-LB053 | GGCGCGGCGAGCGGA | |
| pLB004 | In-Fusion | 4 + 6 | P-LB051 | GCGCGAGGGCGGTGAG | pMK149 |
| | | | P-LB009 | CGGGGTCGCCATCACCGaGGCACCGCCGACGACC | |
| | | 2 + 8 | P-LB008 | GGTCGTCGGCGGTGCCtCGGTGATGGCGACCCCG | pMK149 |
| | | | P-LB050 | TCCGCTCGCCGCGCC | |
| | | pLB003V | P-LB052 | CTCACCGCCCTCGCGC | pMK149 |
| | | | P-LB053 | GGCGCGGCGAGCGGA | |
| pLB005 | In-Fusion | 11 + 17 + 21 + 35 | P-LB050 | TCCGCTCGCCGCGCC | pMK149 |
| | | | P-LB051 | GCGCGAGGGCGGTGAG | |
| | | pLB003V | P-LB052 | CTCACCGCCCTCGCGC | pMK149 |
| | | | P-LB053 | GGCGCGGCGAGCGGA | |
| pLB006 | In-Fusion | 22 + 18 | P-LB051 | GCGCGAGGGCGGTGAG | pMK149 |
| | | | P-LB019 | CGGGGTCGCCATCACCGtGGCACCGCCGACGACC | |
| | | 12 + 36 | P-LB018 | GGTCGTCGGCGGTGCCaCGGTGATGGCGACCCCG | pMK149 |
| | | | P-LB050 | TCCGCTCGCCGCGCC | |
| | | pLB003V | P-LB052 | CTCACCGCCCTCGCGC | pMK149 |
| | | | P-LB053 | GGCGCGGCGAGCGGA | |

| Plasmid | Cloning Method | Fragments | Primer Name | Primer Sequence 5'-3' | Template |
|---|---|---|---|---|---|
| pLB007 | In-Fusion | pLB007_gblock | | | |
| | | pLB007V | P-LB062opt | TCCGAGGAATACGCCGGTCGCGGTACCGCGC | pMK149 |
| | | | P-LB063opt | TTCGGTATTAGCGGGACGAATGCGCACGTGATCGTC | |
| pLB008 | In-Fusion | 25 + 27 | P-LB051 | GCGCGAGGGCGGTGAG | pMK149 |
| | | | P-LB011 | GACGCCTTCGGAGAAGCCtAACCCGTCGGCGCCGG | |
| | | 13 + 32 | P-LB010 | CCGGCGCCGACGGGTTaGGCTTCTCCGAAGGCGTC | pMK149 |
| | | | P-LB050 | TCCGCTCGCCGCGCC | |
| | | pLB003V | P-LB052 | CTCACCGCCCTCGCGC | pMK149 |
| | | | P-LB053 | GGCGCGGCGAGCGGA | |
| pLB009 | In-Fusion | pLB009gblock | | | |
| | | pLB009V | P-MK550 | CCCCAAGAATACGCCGGTCGCGGTACCGCGC | pMK149 |
| | | | P-MK551 | CGCGCACCCTCGGGGCCCGCGCAGCGCAGGG | |
| pLB010 | In-Fusion | pLB010gblock | | | |
| | | pLB010V | P-MK552 | CGCCACACCAAGGAAGACGCCGGTCGCGGTACC | pMK149 |
| | | | P-MK553 | TTCGGGATCTCTGGGACGAATGCGCACGTGATCGTC | |
| pLB011 | In-Fusion | pLB011gblock | | | |
| | | pLB011V | P-MK554 | TCCTAAGAAGACGCCGGTCGCGGTACCGCGC | pMK149 |
| | | | P-MK555 | CCTGCCCCTTCAGGGCCCGCGCAGCGCAGG | |
| pLB013 | In-Fusion | 16 + 32 | P-LB008 | GGTCGTCGGCGGTGCCtCGGTGATGGCGACCCCG | pMK149 |
| | | | P-LB050 | TCCGCTCGCCGCGCC | |
| | | 22 + 19 | P-LB050 | TCCGCTCGCCGCGCC | pMK149 |
| | | | P-LB009 | CGGGGTCGCCATCACCGaGGCACCGCCGACGACC | |
| | | pLB003V | P-LB052 | CTCACCGCCCTCGCGC | pMK149 |
| | | | P-LB053 | GGCGCGGCGAGCGGA | |

| Plasmid | Cloning Method | Fragments | Primer Name | Primer Sequence 5'-3' | Template |
|---|---|---|---|---|---|
| | | | P-LB015 | GGGTGACGCCTTCGGAGtAGCCtAACCCGTCGGCGCCGG | |
| pLB015 | In-Fusion | 21 | | | |
| | | 30 + 37 | P-LB016 | GCGTCTTCCTCGGAGTGGaGAAGTaCGGCTACGGCGAGGAC | pMK149 |
| | | | P-LB050 | TCCGCTCGCCGCGCC | |
| | | pLB003V | P-LB052 | CTCACCGCCCTCGCGC | pMK149 |
| | | | P-LB053 | GGCGCGGCGAGCGGA | |
| pLB016 | In-Fusion | pLB016gblock | | | |
| | | pLB016V | P-LB073 | GGCCCCCAAGAAGACGCCGGTGTCGC | pMK147 |
| | | | P-LB074 | GCGCCTTCGGGCGTCGCCCAGCAGC | |
| pLB017 | In-Fusion | pLB017gblock | | | |
| | | pLB016V | P-LB073 | GGCCCCCAAGAAGACGCCGGTGTCGC | pMK147 |
| | | | P-LB074 | GCGCCTTCGGGCGTCGCCCAGCAGC | |
| pLB018 | In-Fusion | pLB018gblock | | | |
| | | pLB016V | P-LB073 | GGCCCCCAAGAAGACGCCGGTGTCGC | pMK147 |
| | | | P-LB074 | GCGCCTTCGGGCGTCGCCCAGCAGC | |
| pLB019 | In-Fusion | pLB019gblock | | | |
| | | pLB016V | P-LB073 | GGCCCCCAAGAAGACGCCGGTGTCGC | pMK147 |
| | | | P-LB074 | GCGCCTTCGGGCGTCGCCCAGCAGC | |
| pLB021 | In-Fusion | pLB021gblock | | | |
| | | pLB016V | P-LB073 | GGCCCCCAAGAAGACGCCGGTGTCGC | pMK147 |
| | | | P-LB074 | GCGCCTTCGGGCGTCGCCCAGCAGC | |
| pLB022 | In-Fusion | pLB022gblock | | | |
| | | pLB016V | P-LB073 | GGCCCCCAAGAAGACGCCGGTGTCGC | pMK147 |
| | | | P-LB074 | GCGCCTTCGGGCGTCGCCCAGCAGC | |

| Plasmid | Cloning Method | Fragments | Primer Name | Primer Sequence 5'-3' | Template |
|---|---|---|---|---|---|
| pLB023 | In-Fusion | pLB023gblock | | | |
| | | pLB016V | P-LB073 | GGCCCCCAAGAAGACGCCGGTGTCGC | pMK147 |
| | | | P-LB074 | GCGCCTTCGGGCGTCGCCCAGCAGC | |
| pLB024 | In-Fusion | pLB024gblock | | | |
| | | pLB024V | P-LB073 | GGCCCCCAAGAAGACGCCGGTGTCGC | pMK147 |
| | | | P-LB074 | GCGCCTTCGGGCGTCGCCCAGCAGC | |
| pLB026 | In-Fusion | pLB026gblock | | | |
| | | pLB016V | P-LB073 | GGCCCCCAAGAAGACGCCGGTGTCGC | pMK147 |
| | | | P-LB074 | GCGCCTTCGGGCGTCGCCCAGCAGC | |
| pLB027 | In-Fusion | pLB027glbock | | | |
| | | pLB016V | P-LB073 | GGCCCCCAAGAAGACGCCGGTGTCGC | pMK147 |
| | | | P-LB074 | GCGCCTTCGGGCGTCGCCCAGCAGC | |
| pMK178 | In-Fusion | pMK178_I | P-MK517 | GAGCGCGTCTGGCTCCTGCCGGACCGCACCAC | pMK175 |
| | | | P-MK518 | GCCACCGGATCCGCCGCCGAGCTCACTAGTGAGG | |
| | | pMK178_V | P-MK312 | GGATCCGCCACCGGATCCGCCTTCAGCAACATCGTTCTCCAATCTG | pMK111 |
| | | | P-MK483 | GGCGGATCCGGTGGCGGA | |
| pMK179 | In-Fusion | pMK179_I | P-MK519 | GAGCGCGTCTGGCTCCCCATCCCCACCGGCG | pMK168 |
| | | | P-MK520 | GCCACCGGATCCGCCGACGAGCCGCTCCAGGTA | |
| | | pMK179_V | P-MK312 | GGATCCGCCACCGGATCCGCCTTCAGCAACATCGTTCTCCAATCTG | pMK111 |
| | | | P-MK483 | GGCGGATCCGGTGGCGGA | |
| pMK183 | In-Fusion | pMK183_I | P-MK260 | GAAGGAGATATACATATGAGCGGTGACAACGGCATGACCGAGGAAAAG | pMK178 |

| Plasmid | Cloning Method | Fragments | Primer Name | Primer Sequence 5'-3' | Template |
|---|---|---|---|---|---|
| | | | P-MK560 | CCGGTCGCGCAGGCTC | |
| | | pMK183_V | P-MK559 | ACGGAGAGCCTGCGCGACCGGCTGGCGTCGCTGCCCG | pMK150 |
| | | | P-MK239 | CATATGTATATCTCCTTCTTAAAGTTAAACAAAATTA | |
| pMK184 | In-Fusion | pMK184_I | P-MK260 | GAAGGAGATATACATATGAGCGGTGACAACGGCATGACCGAGGAAAAG | pMK150 |
| | | | P-MK562 | CGCGCCCACCCGCGG | |
| | | pMK184_V | P-MK239 | CATATGTATATCTCCTTCTTAAAGTTAAACAAAATTA | pMK178 |
| | | | P-MK561 | GCCGAACCGCGGGTGGGCGCGCTGGCGGGTCTGCCGC | |
| pMK185 | In-Fusion | pMK185_I | P-MK260 | GAAGGAGATATACATATGAGCGGTGACAACGGCATGACCGAGGAAAAG | pMK150 |
| | | | P-MK562 | CGCGCCCACCCGCGG | |
| | | pMK185_V | P-MK563 | GCCGAACCGCGGGTGGGCGCGCTCGCGGCGCTGTCGAC | pMK179 |
| | | | P-MK239 | CATATGTATATCTCCTTCTTAAAGTTAAACAAAATTA | |
| pLB001 | QuickChange | - | P-LB002 | GGCGTCTTCCTCGGAGTGtgGAAGTTCGGCTACGGCGAG | pMK149 |
| | | | P-LB003 | CTCGCCGTAGCCGAACTTCcaCACTCCGAGGAAGACGCC | |
| pLB002 | QuickChange | - | P-LB004 | CGTCTTCCTCGGCGCCtgGTACCAGGGCTACGGCC | pMK147 |
| | | | P-LB005 | GGCCGTAGCCCTGGTACcaGGCGCCGAGGAAGACG | |

**Table S8. Cloning strategy of fragments used in In-Fusion cloning and primer sequences**

| Fragment | Primer Name | Primer Sequence 5'-3' | Template | Used to generate |
|---|---|---|---|---|
| 1 | P-LB008 | GGTCGTCGGCGGTGCCtCGGTGATGGCGACCCCG | pMK149 | pLB003 |
| | P-LB011 | GACGCCTTCGGAGAAGCCtAACCCGTCGGCGCCGG | | |
| 2 | P-LB008 | GGTCGTCGGCGGTGCCtCGGTGATGGCGACCCCG | pMK149 | pLB004 |
| | P-LB015 | GGGTGACGCCTTCGGAGtAGCCtAACCCGTCGGCGCCGG | | |
| 3 | P-LB006 | GGCGTCTTCCTCGGAGTGcaGAAGTaCGGCTACGGCGAGGAC | pMK149 | pLB003 |
| | P-LB009 | CGGGGTCGCCATCACCGaGGCACCGCCGACGACC | | |
| 4 | P-LB012 | GGCGTCTTCCTCGGAGTGctGAAGTaCGGCTACGGCGAGGAC | pMK149 | pLB004 |
| | P-LB009 | CGGGGTCGCCATCACCGaGGCACCGCCGACGACC | | |
| 5 | P-LB051 | GCGCGAGGGCGGTGAG | pMK149 | pLB003 |
| | P-LB007 | GTCCTCGCCGTAGCCGtACTTCtgCACTCCGAGGAAGACGCC | | |
| 6 | P-LB051 | GCGCGAGGGCGGTGAG | pMK149 | pLB004 |
| | P-LB013 | GTCCTCGCCGTAGCCGtACTTCagCACTCCGAGGAAGACGCC | | |
| 7 | P-LB010 | CCGGCGCCGACGGGTTaGGCTTCTCCGAAGGCGTC | pMK149 | pLB003 |
| | P-LB050 | TCCGCTCGCCGCGCC | | |
| 8 | P-LB014 | CCGGCGCCGACGGGTTaGGCTaCTCCGAAGGCGTCACCC | pMK149 | pLB004 |
| | P-LB050 | TCCGCTCGCCGCGCC | | |
| 11 | P-LB018 | GGTCGTCGGCGGTGCCaCGGTGATGGCGACCCCG | pMK149 | pLB005 |
| | P-LB021 | CAGGGTGACGCCTTCGGAccAGCCGtACCCGTCGGCGCCGG | | |
| 12 | P-LB018 | GGTCGTCGGCGGTGCCaCGGTGATGGCGACCCCG | pMK149 | pLB006 |
| | P-LB023 | CAGGGTGACGCCTTCGGAcAtGCCtAACCCGTCGGCGCCGG | | |
| 13 | P-LB010 | CCGGCGCCGACGGGTTaGGCTTCTCCGAAGGCGTC | pMK149 | pLB008 |
| | P-LB035 | CCCGCTCGGCGCGGgAAGCCCGTTGCTGGCCC | | |
| 16 | P-LB008 | GGTCGTCGGCGGTGCCtCGGTGATGGCGACCCCG | pMK149 | pLB013 |
| | P-LB035 | CCCGCTCGGCGCGGgAAGCCCGTTGCTGGCCC | | |
| 17 | P-LB016 | GCGTCTTCCTCGGAGTGGaGAAGTaCGGCTACGGCGAGGAC | pMK149 | pLB005 |
| | P-LB019 | CGGGGTCGCCATCACCGtGGCACCGCCGACGACC | | |

| Fragment | Primer Name | Primer Sequence 5'-3' | Template | Used to generate |
|---|---|---|---|---|
| 18 | P-LB006 | GGCGTCTTCCTCGGAGTGcaGAAGTaCGGCTACGGCGAGGAC | pMK149 | pLB006 |
| | P-LB019 | CGGGGTCGCCATCACCGtGGCACCGCCGACGACC | | |
| 19 | P-LB006 | GGCGTCTTCCTCGGAGTGcaGAAGTaCGGCTACGGCGAGGAC | pMK149 | pLB013 |
| | P-LB009 | CGGGGTCGCCATCACCGaGGCACCGCCGACGACC | | |
| 21 | P-LB051 | GCGCGAGGGCGGTGAG | pMK149 | pLB015 |
| | P-LB017 | GTCCTCGCCGTAGCCGtACTTCtCCACTCCGAGGAAGACGC | | |
| 22 | P-LB051 | GCGCGAGGGCGGTGAG | pMK149 | pLB006 |
| | P-LB007 | GTCCTCGCCGTAGCCGtACTTCtgCACTCCGAGGAAGACGCC | | |
| 25 | P-LB051 | GCGCGAGGGCGGTGAG | pMK149 | pLB008 |
| | P-LB033 | GTGTCCTCGCCGTAGCCctgCTTCGtCACTCCGAGGAAGAC | | |
| 27 | P-LB032 | GTCTTCCTCGGAGTGaCGAAGcagGGCTACGGCGAGGACAC | pMK149 | pLB008 |
| | P-LB011 | GACGCCTTCGGAGAAGCCtAACCCGTCGGCGCCGG | | |
| 30 | P-LB016 | GCGTCTTCCTCGGAGTGGaGAAGTaCGGCTACGGCGAGGAC | pMK149 | pLB015 |
| | P-LB015 | GGGTGACGCCTTCGGAGtAGCCtAACCCGTCGGCGCCGG | | |
| 32 | P-LB034 | GGGCCAGCAACGGGCTTcCCGCGCCGAGCGGG | pMK149 | pLB008 |
| | P-LB050 | TCCGCTCGCCGCGCC | | |
| 35 | P-LB020 | CCGGCGCCGACGGGTaCGGCTggTCCGAAGGCGTCACCCTG | pMK149 | pLB005 |
| | P-LB050 | TCCGCTCGCCGCGCC | | |
| 36 | P-LB022 | CCGGCGCCGACGGGTTaGGCaTgTCCGAAGGCGTCACCCTG | pMK149 | pLB006 |
| | P-LB050 | TCCGCTCGCCGCGCC | | |
| 37 | P-LB014 | CCGGCGCCGACGGGTTaGGCTaCTCCGAAGGCGTCACCC | pMK149 | pLB015 |
| | P-LB050 | TCCGCTCGCCGCGCC | | |

**Table S9. gBlocks used for In-Fusion cloning**

| gblock | Sequence 5'-3' |
|---|---|
| pLB007_ gblock | GGCGTATTCCTCGGAGTGCAGAAAAAGGATATGGCGAAGACACGGCTGCAGCTGAGGACGTCGAGGGATACAGTGTTACCGGCGTGGCGCCTGCCGTGGCTAGTGGT CGAATATCGTACACGATGGGTCTGGAGGGTCCAAGTATTTCCGTAGATACGGCCTGTAGTTCGTCCCTGGTAGCACTGCATTTGGCGGTAGAGTCGCTGCGAAAAGGT GAAAGTTCCATGGCCGTGGTAGGTGGTGCTGCGGTAATGGCAACACCTGGAGTATTCGTGGACTTTTCCCGTCAACGTGCGTTGGCAGCCGACGGTCGTAGTAAGGCG TTTGGCGCGGGAGCGGATGGTTGGGGATTTAGTGAAGGCGTGACCCTGGTGCTCCTCGAGCGACTGAGCGAAGCTCGACGCAATGGTCATGAGGTTTTGGCAGTGGTC CGCGGTTCGGCGCTCAATCAAGACGGAGCAAGCAATGGTTTGACTGCTCCTAGCGGCCCTGCACAAAGGCGCGTGATCCGGCAGGCGCTGGAGTCCTGCGGTCTGGAG CCGGGCGACGTTGATGCTGTCGAAGCTCACGGTACGGGTACTGCGTTGGGTGATCCGATTGAGGCGAATGCGCTCCTCGATACCTATGGACGAGATCGTGATGCCGAC CGACCGTTGTGGCTGGGATCGGTTAAGTCCAACATTGGACATACTCAAGCTGCGGCCGGTGTTACTGGTTTGCTGAAGGTAGTACTCGCACTCCGTAATGGCGAACTG CCAGCAACCCTCCATGTAGAGGAGCCGACACCACATGTAGATTGGTCGTCCGGAGGTGTTGCGCTCCTGGCCGGAAATCAACCATGGAGGAGGGGAGAGCGAACGCGA CGGGCGAGGGTATCGAGCTTCGGTATTAGCGGG |
| pLB009_ gblock | GGCGTATTCTTGGGGGTGCAGAAGAACGGCTATGGCGAAGATACTGCGGCCGCAGAAGATGTCGAGGGTTACTCAGTTACCGGAGTCGCGCCCGCCGTAGCGTCTGGG CGCATCTCTTATACCATGGGCCTTGAGGGACCCTCAATCAGTGTTGACACCGCTTGTTCATCAAGCCTGGTCGCGTTGCATTTAGCCGTCGAATCATTACGTAAGGGG GAGTCCAGCATGGCAGTTGTGGGGGGAGCAGCGGTGATGGCCACGCCGGGTGTTTTTGTGGATTTTTCGCGCCAACGTGCACTTGCCGCCGATGGGCGTTCAAAGGCG TTCGGTGCAGGCGCGGATGGGTTAGGATTTAGTGAGGGTGTGACCCTTGTTCTTTTGGAGCGTTTGAGTGAGGCTCGCCGTAACGGACATGAGGTGTTGGCGGTTGTA CGTGGTAGCGCGTTAAATCAAGACGGCGCAAGTAACGGTTTACGCGCACCCTCGGGGC |
| pLB010_ gblock | CTTCCTTGGTGTGGCGAAATATGGCTATGGAGAGGACACCGCTGCGGCAGAAGATGTGGAGGGTTACTCTGTCACAGGAGTGGCACCTGCGGTCGCTTCCGGGCGTAT TTCCTACACAATGGGGTTAGAAGGTCCCAGTATCTCGGTAGACACTGCATGCTCGTCTTCTCTGGTCGCGTTGCATTTGGCTGTGGAGAGTCTTCGCAAAGGGGAGTC AAGTATGGCTGTAGTCGGTGGGGCCACAGTAATGGCAACGCCCGGGGTCTTTGTAGACTTTTCTCGCCAGCGCGCACTTGCCGCGGACGGACGTTCCAAGGCGTTTGG TGCCGGGGCCGACGGGGCTGGGTTTTCTGAAGGGGTAACATTAGTGTTGCTTGAGCGTTTATCAGAAGCCCGTCGTAATGGACATGAAGTACTTGCGGTAGTCCGCGG TAGTGCGCTGAACCAAGATGGAGCCAGCAACGGTCTGACTGCCCCCTCGGGACCCGCTCAGCGCCGTGTGATTCGCCAAGCGCTGGAATCTTGCGGTTTAGAGCCCGG GGATGTCGACGCCGTAGAAGCCCATGGCACTGGAACAGCTTTGGGTGACCCAATCGAGGCTAACGCTCTTCTTGACACTTATGGTCGCGATCGTGACGCCGATCGTCC GCTGTGGTTAGGTTCGGTCAAATCAAACATTGGCCACACGCAGGCTGCAGCAGGCGTAACGGGACTTTTGAAAGTAGTGCTGGCGCTTCGTAATGGGGAGTTACCAGC TACTTTGCACGTTGAGGAACCCACTCCTCATGTAGATTGGTCGTCGGGCGGCGTTGCCTTACTGGCGGGTAACCAACCGTGGCGCCGTGGTGAACGTACTCGTCGTGC GCGTGTTAGTTCGTTCGGGATCTCTGGG |
| pLB011_ gblock | GGCGTCTTCTTAGGAGTCCAGAAGTATGGCTATGGGGAGGACACCGCTGCTGCGGAGGACGTAGAAGGTTATTCTGTCACAGGTGTAGCGCCTGCCGTGGCGTCAGGC CGCATCTCTTACACAATGGGGTTAGAAGGTCCTTCAATTAGCGTAGACACCGCTTGCTCATCGTCGTTGGTAGCTCTTCACCTTGCAGTAGAGTCTTTACGCAAGGGG GAGTCTTCCATGGCCGTAGTGGGTGGTGCCTGCGTAATGGCAACCCCTGGCGTGTTCGTAGATTTCAGCCGCCAACGTGCATTAGCAGCAGATGGGCGCAGCAAAGCA TTTGGGGCTGGTGCGGATGGATTAGGTTTTTCCGAAGGGGTTACTTTAGTTCTTTTGGAGCGTTTGTCAGAGGCGCGTCGTAATGGTCATGAGGTGTTAGCAGTGGTT CGCGGCTCTGCCCTGAATCAAGACGGGGCAAGTAACGGCTTGCCTGCCCCTTCAGGG |
| pLB016_ gblock | GTCTTCTTGGGGGCCAGCTATCATGGTTATGGCCAAGATGCAGTGGTACCTGAGGATTCTGAGGGATATCTGCTTACGGGAAACTCGTCTGCGGTTGTAAGTGGGCGT GTTGCTTATGTACTGGGCTTGGAAGGGCCTGCGGTTACGGTAGATACAGCATGTTCCGCATCTTTGGTAGCTTTGCACAGCGCATGCGGATCTCTTCGTGATGGTGAT TGCGGTCTTGCCGTAGCAGGCGGAGTTAGCGTAATGGCCGGTCCAGAAATCTTTACAGAGTTCTCGCGCCAGGGCGGCCTGGCCGTCGATGGGCGTTGTAAGGCCTTT AGCGCAGAGGCGGATGGTTTCGGTTTCGCTGAAGGGGTAGCTGTGGTCTTGTTACAACGCCTTAGTGATGCTCGCCGTGCGGGCCGCCAGGTCTTAGGGGTTGTAGCG GGAAGTGCAATTAATCAAGATGGTGCATCAAACGGACTGTCGGCGCCTTCGGGCGTC |
| pLB017_ gblock | GTCTTTTTGGGAGCTTGGTATAGTGGTTATGGGCAAGATGCCGTTGTCCCTGAAGACAGCGAAGGATACTTGCTTACAGGGAACAGTTCCGCCGTCGTGTCGGGCCGC GTTGCTTACGTGTTGGGTTTAGAAGGGCCCGCTGTAACGGTAGATACAGCTTGTTCCGCTAGCCTGGTAGCTTTACATTCCGCCTGTGGCTCACTGCGCGATGGTGAT TGTGGATTAGCAGTGGCAGGAGGGGTGGCGGTAATGGCTGGACCCGAGGTTTTTACCGAATTTTCTCGTCAGGGTGGGTTAGCTGTTGACGGTCGCTGTAAGGCGTTT AGCGCCGAGGCCGATGGGTTCGGCTTTGCCGAGGGTGTTGCAGTAGTTCTTCTTCAGCGTTTGTCCGATGCCCGCCGTGCAGGTCGTCAAGTGCTGGGAGTTGTCGCG GGTTCTGCGATTAACCAAGATGGCGCGAGCAACGGACTTCGCGCCCCATCGGGCGTC |
| pLB018_ gblock | GTCTTTCTGGGGGCATGGTATCATGGGTACGGCCAAGATGCCGTGGTGCCAGAGGACTCGGAGGGCTATTTATTAACAGGTAATTCTTCAGCGGTAGTGAGTGGGCGT GTCGCCTACGTTCTGGGGTTAGAGGGGCCAGCTGTCACGGTGGATACAGCATGCAGTGCTTCGTTGGTCGCTTTACATTCGGCCTGTGGTTCTTTGCGTGATGGTGAT TGCGGTCTGGCAGTAGCAGGAGGGGTAACTGTCATGGCCGGTCCTGAAGTGTTTACTGAATTCTCTCGTCAAGGAGGGTTGGCCGTAGATGGCCGCTGTAAGGCATTT TCCGCTGAAGCAGACGGGTTCGGGTTCGCAGAAGGAGTGGCAGTCGTTTTGCTGCAGCGCTTATCGGATGCTCGTCGCGCTGGACGCCAGGTGTTAGGAGTGGTGGCG |

| gblock | Sequence 5'-3' |
|---|---|
| | GGCTCCGCGATCAATCAGGATGGCGCTAGCAACGGCCTTACAGCCCCTTCTGGAGTC |
| pLB019_gblock | GTCTTTTTAGGTGCCTCATATCATGGATATGGCCAGGATGCTGTCGTTCCCGAGGATTCCGAGGGCTATCTGCTTACGGGTAACTCTTCTGCGGTAGTAAGCGGACGT<br>GTAGCTTATGTCCTTGGCTTAGAAGGACCTGCTGTGACTGTCGACACAGCGTGCTCCGCCAGCTTAGTTGCGCTTCATAGTGCGTGTGGTAGTTTACGCGACGGAGAC<br>TGTGGTCTGGCCGTCGCGGGAGGAGTTCATGTAATGGCAGGCCCAGAGGTGTTTACTGAGTTTAGCCGTCAAGGAGGTTTAGCGGTAGACGGACGTTGCAAGGCCTTC<br>AGCGCCGAGGCTGACGGGTTCGGATTTGCAGAAGGGGTAGCTGTGGTACTGCTGCAACGCTTGTCAGACGCCCGTCGCGCAGGACGTCAGGTATTGGGAGTAGTTGCT<br>GGGTCTGCTATTAACCAGGATGGCGCGTCAAACGGGTTATCTGCCCCGTCGGGGGTC |
| pLB021_gblock | GTCTTTTTAGGCGCGTGGTACTCAGGTTACGGCCAGGACGCTGTTGTGCCCGAAGATTCAGAGGGTTACCTTTTAACCGGGAACTCCAGTGCAGTCGTAAGTGGTCGC<br>GTAGCCTACGTTTTAGGTTTAGAGGGTCCCGCCGTCACTGTGGATACGGCGTGTAGCAGTTCCTTAGTAGCGTTGCACAGTGCTTGCGGATCATTACGCGATGGTGAC<br>TGTGGCCTGGCAGTAGCGGGTGGTGTAGCCGTTATGGCGGGCCCAGAAGTTTTCACCGAATTTTCTCGTCAAGGGGGTTTAGCTGTTGACGGCCGCTGTAAGGCTTTC<br>TCTGCAGAGGCTGATGGTTTTGGCTTTGCAGAGGGGGTAGCCGTAGTCCTGCTGCAGCGTTTATCGGACGCCCGTCGCGCTGGCCGCCAGGTTCTGGGCGTAGTTGCA<br>GGTTCAGCGATCAATCAAGATGGCGCAAGTAACGGCCTTACTGCCCCTTCCGGCGTC |
| pLB022_gblock | GTCTTCTTGGGGGCGGCGTACTCTGGATATGGTCAAGATGCGGTGGTTCCCGAGGACTCTGAGGGGTACTTGTTGACTGGCAATAGTAGCGCTGTAGTCAGCGGTCGT<br>GTCGCGTACGTGCTTGGTCTTGAGGGCCCAGCCGTAACTGTCGATACTGCATGTTCAGCCAGCCTTGTGGCCCTGCATTCGGCATGCGGATCCTTACGTGACGGAGAC<br>TGCGGACTTGCGGTGGCTGGTGGGGTTTCGGTAATGGCAGGCCCGGAACTTTTTACGGAGTTTTCGCGCCAAGGTGGCTTAGCAGTAGATGGCCGTTGTAAGGCATTT<br>AGTGCCGAAGCGGATGGCTTTGGAATGGCCGAAGGGGTCGCTGTCGTCCTGCTTCAACGTTTGTCGGACGCTCGTCGTGCGGGTCGCCAGGTACTTGGAGTAGTTGCC<br>GGATCAGCAATCAACCAGGATGGGGCTTCAAACGGGTTAGAAGCACCGTCCGGAGTC |
| pLB023_gblock | GTCTTTCTTGGCGCGGGGTATCACGGGTACGGACAGGACGCAGTAGTTCCCGAAGATAGCGAAGGTTACCTGTTGACCGGAAACAGCAGTGCCGTGGTCAGCGGACGT<br>GTCGCGTATGTTCTTGGTTTAGAAGGACCTGCGGTGACTGTTGATACTGCCTGTAGCGCAAGTCTTGTTGCTTTACATAGTGCCTGTGGTTCTTTACGTGATGGAGAC<br>TGTGGGCTGGCGGTGGCCGGAGGCGTGCATGTCATGGCGGGGCCCGAAGTGTTTACAGAGTTCTCACGCCAAGGAGGCCTTGCGGTAGACGGTCGCTGTAAGGCTTTC<br>TCGGCTGAGGCGGATGGTTTCGGCTTTGCGGAAGGGGTCGCCGTTGTATTACTTCAGCGTCTTTCAGATGCTCGTCGCGCCGGTCGTCAGGTCTTGGGCGTGGTAGCG<br>GGATCTGCAATTAATCAGGATGGAGCTAGCAATGGTTTACAGGCTCCGAGCGGCGTC |
| pLB024_gblock | GTCTTCCTTGGGGCGTGGTATCAAGGTTACGGTCAGGACGCCGTAGTTCCTGAAGATTCCGAAGGCTATTTGTTAACGGGGAATTCGTCTGCGGTGGTTTCTGGTCGT<br>GTAGCTTACGTCTTAGGGCTTGAGGGACCTGCTGTCACAGTCGATACAGCTTGTAGCGCATCTTTAGTTGCACTGCACTCGGCATGTGGCTCTCTTCGTGATGGAGAC<br>TGCGGATTAGCAGTTGCGGGGGGCGTCACCGTAATGGCCGGCCCTGAAGTCTTCACCGAATTTTCGCGTCAAGGAGGTCTGGCAGTCGACGGACGTTGTAAGGCATTT<br>AGTGCAGAAGCCGATGGTTTTGGGTTCGCCGAAGGTGTCGCCGTTGTGCTGCTGCAGCGTCTGTCAGACGCGCGTCGCGCCGGACGCCAGGTATTGGGTGTGGTAGCG<br>GGAAGTGCGATCAACCAAGATGGCGCGTCTAATGGCTTGAGTGCCCCCAGTGGGGTGGCGCAACAGCGTGTGATCCGTAAAGCCTGGGCACGCGCAGGAATTACTGGA<br>GCGGATGTAGCCGTGGTAGAGGCGCACGGTACAGGGACACGTCTGGGAGACCCAGTCGAAGCGTCCGCTTTGTTAGCGACGTACGGAAAATCACGTGGCTCTAGTGGT<br>CCCGTACTTTTGGGCTCGGTTAAGTCTAACATTGGGCACGCCCAAGCTGCAGCGGGAGTGGCGGGCGTAATTAAGGTTGTCTTAGGGTTAAATCGCGGTTTGGTCCCG<br>CCCATGTTATGTCGCGGTGAACGCAGTCCACTGATTGAATGGAGCTCCGGCGGTGTGGAGCTGGCTGAAGCTGTCAGCCCCTGGCCACCTGCCGCTGACGGAGTGCGT<br>CGTGCTGGAGTAAGCTCTTTTGGGGTTTCGGGG |
| pLB026_gblock | GTCTTTTTGGGCGCCGCATACCAAGGTTACGGTCAGGATGCCGTTGTTCCCGAAGACTCCGAGGGTTACCTGCTTACTGGTAATAGCAGTGCGGTGGTCTCAGGTCGC<br>GTAGCATACGTGTTGGGTCTGGAGGGGCCTGCCGTCACAGTCGACACGGCTTGCAGTGCGAGTTTAGTTGCTTTACACAGTGCTTGTGGCAGTCTTCGTGATGGTGAT<br>TGCGGATTAGCCGTTGCTGGTGGTGTGTCTGTGATGGCTGGGCCGGAGCTTTTTACTGAGTTTAGCCGTCAAGGGGGCCTGGCAGTAGATGGACGTTGCAAGGCGTTT<br>TCCGCCGAGGCAGACGGGTTCGGAATGGCTGAGGGGGTGGCGGTGGTCCTTCTTCAACGCCTTTCAGATGCGCGTCGTGCAGGGCGTCAAGTTTTAGGGGTAGTCGCC<br>GGTTCTGCAATTAACCAGGATGGGGCCAGCAACGGGTTAGCGGCACCTTCTGGGGTC |
| pLB027_gblock | GTCTTTCTTGGGGCGACTTACCAGGGGTACGGTCAAGACGCGGTTGTGCCCGAGGACTCGGAAGGTTACTTGTTAACCGGAAATAGTTCAGCGGTCGTGAGTGGGCGC<br>GTGGCCTATGTCTTGGGCTTGGAAGGCCCGGCAGTGACCGTGGACACAGCTTGCAGCGCAAGTTTAGTCGCTCTGCATAGTGCCTGTGGATCTTTACGCGATGGCGAT<br>TGCGGCTTGGCTGTCGCCGGTGGCGTCCATGTTATGGCTGGCCCAGAGATGTTTACTGAGTTCTCTCGCCAGGGAGGCTTGGCAGTTGATGGGCGCTGCAAAGCATTC<br>TCTGCGGAGGCGGACGGTTTTGGATTCGCGGAGGGGGTAGCTGTAGTATTGTTACAACGTTTGAGCGATGCCCGTCGTGCTGGCCGTCAGGTTCTTGGTGTAGTGGCC<br>GGTTCAGCTATCAACCAAGACGGTGCATCCAATGGTTTGTCCGCCCCAAGTGGCGTC |

**Table S10. Amino acid sequences of newly generated substrate-donating modules**

| Construct | Amino acid sequence |
|---|---|
| MK178 (5)KS1-AT1-KR2-ACP2-SZ3 | MSGDNGMTEEKLRRYLKRTVTELDSVTARLREVEHRAGEPVAVVAMACRLPGGVSTPEEFWELLSEGRDAVAGLPTDRGWDLDSLFH<br>PDPTRSGTAHQRGGGFLTEATAFDPAFFGMSPREALAVDPQQRLMLELSWEVLERAGIPPTSLQASPTGVFVGLIPQEYGPRLAEGG<br>EGVEGYLMTGTTTSVASGRIAYTLGLEGPAISVDTACSSSLVAVHLACQSLRRGESSLAMAGGVTVMPTPGMLVDFSRMNSLAPDGR<br>CKAFSAGANGFGMAEGAGMLLLERLSDARRNGHPVLAVLRGTAVNSDGASNGLSAPNGRAQVRVIQQALAESGLGPADIDAVEAHGT<br>GTRLGDPIEARALFEAYGRDREQPLHLGSVKSNLGHTQAAAGVAGVIKMVLAMRAGTLPRTLHASERSKEIDWSSGAISLLDEPEPW<br>PAGARPRRAGVSSFGISGTNAHAIIEEAPQVVEGERVEAGDVVAPWVLSASSAEGLRAQAARLAAHLREHPGQDPRDIAYSLATGRA<br>ALPHRAAFAPVDESAALRVLDGLATGNADGAAVGTSRAQQRAVFVFPGQGWQWAGMAVDLLDTSPVFAAALRECADALEPHLDFEVI<br>PFLRAEAARREQDAALSTERVDVVQPVMFAVMVSLASMWRAHGVEPAAVIGHSQGEIAAACVAGALSLDDAARVVALRSRVIATMPG<br>NKGMASIAAPAGEVRARIGDRVEIAAVNGPRSVVVAGDSDELDRLVASCTTECIRAKRLAVDYASHSSHVETIRDALHAELGEDFHP<br>LPGFVPFFSTVTGRWTQPDELDAGYWYRNLRRTVRFADAVRALAEQGYRTFLEVSAHPILTAAIEEIGDGSGADLSAIHSLRRGDGS<br>LADFGEALSRAFAAGVAVDWESVHLGTGARRVPLPTYPFQRERVWLLPDRTTPRDELDGWFYRVDWTEVPRSEPAALRGRWLVVVPE<br>GHEEDGWTVEVRSALAEAGAEPEVTRGVGGLVGDCAGVVSLLALEGDGAVQTLVLVRELDAEGIDAPLWTVTFGAVDAGSPVARPDQ<br>AKLWGLGQVASLERGPRWTGLVDLPHMPDPELRGRLTAVLAGSEDQVAVRADAVRARRLSPAHVTATSEYAVPGGTILVTGGTAGLG<br>AEVARWLAGRGAEHLALVSRRGPDTEGVGDLTAELTRLGARVSVHACDVSSREPVRELVHGLIEQGDVVRGVVHAAGLPQQVAINDM<br>DEAAFDEVVAAKAGGAVHLDELCSDAELFLLFSSGAGVWGSARQGAYAAGNAFLDAFARHRRGRGLPATSVAWGLWAAGGMTGDEEA<br>VSFLRERGVRAMPVPRALAALDRVLASGETAVVVTDVDWPAFAESYTAARPRPLLDRIVTTAPSERAGEPETESLRDRLAGLPRAER<br>TAELVRLVRTSTATVLGHDDPKAVRATTPFKELGFDSLAAVRLRNLLNAATGLRLPSTLVFDHPNASAVAGFLTSELGGGSGGGSGN<br>EVTTLENDAAFIENENAYLEKEIARLRKEKAALRNRLAHKKLEHHHHHH |
| MK179 (5)KS1-AT1-KR5-ACP5-SZ3 | MSGDNGMTEEKLRRYLKRTVTELDSVTARLREVEHRAGEPVAVVAMACRLPGGVSTPEEFWELLSEGRDAVAGLPTDRGWDLDSLFH<br>PDPTRSGTAHQRGGGFLTEATAFDPAFFGMSPREALAVDPQQRLMLELSWEVLERAGIPPTSLQASPTGVFVGLIPQEYGPRLAEGG<br>EGVEGYLMTGTTTSVASGRIAYTLGLEGPAISVDTACSSSLVAVHLACQSLRRGESSLAMAGGVTVMPTPGMLVDFSRMNSLAPDGR<br>CKAFSAGANGFGMAEGAGMLLLERLSDARRNGHPVLAVLRGTAVNSDGASNGLSAPNGRAQVRVIQQALAESGLGPADIDAVEAHGT<br>GTRLGDPIEARALFEAYGRDREQPLHLGSVKSNLGHTQAAAGVAGVIKMVLAMRAGTLPRTLHASERSKEIDWSSGAISLLDEPEPW<br>PAGARPRRAGVSSFGISGTNAHAIIEEAPQVVEGERVEAGDVVAPWVLSASSAEGLRAQAARLAAHLREHPGQDPRDIAYSLATGRA<br>ALPHRAAFAPVDESAALRVLDGLATGNADGAAVGTSRAQQRAVFVFPGQGWQWAGMAVDLLDTSPVFAAALRECADALEPHLDFEVI<br>PFLRAEAARREQDAALSTERVDVVQPVMFAVMVSLASMWRAHGVEPAAVIGHSQGEIAAACVAGALSLDDAARVVALRSRVIATMPG<br>NKGMASIAAPAGEVRARIGDRVEIAAVNGPRSVVVAGDSDELDRLVASCTTECIRAKRLAVDYASHSSHVETIRDALHAELGEDFHP<br>LPGFVPFFSTVTGRWTQPDELDAGYWYRNLRRTVRFADAVRALAEQGYRTFLEVSAHPILTAAIEEIGDGSGADLSAIHSLRRGDGS<br>LADFGEALSRAFAAGVAVDWESVHLGTGARRVPLPTYPFQRERVWLPIPTGGRARDEDDDWRYQVVWREAEWESASLAGRVLLVTGP<br>GVPSELSDAIRSGLEQSGATVLTCDVESRSTIGTALEAADTDALSTVVSLLSRDGEAVDPSLDALALVQALGAAGVEAPLWVLTRNA<br>VQVADGELVDPAQAMVGGLGRVVGIEQPGRWGGLVDLVDADAASIRSLAAVLADPRGEEQVAIRADGIKVARLVPAPARAARTRWSP<br>RGTVLVTGGTGGIGAHVARWLARSGAEHLVLLGRRGADAPGASELREELTALGTGVTIAACDVADRARLEAVLAAERAEGRTVSAVM<br>HAAGVSTSTPLDDLTEAEFTEIADVKVRGTVNLDELCPDLDAFVLFSSNAGVWGSPGLASYAAANAFLDGFARRRRSEGAPVTSIAW<br>GLWAGQNMAGDEGGEYLRSQGLRAMDPDRAVEELHITLDHGQTSVSVVDMDRRRFVELFTAARHRPLFDEIAGARAEARQSEEGPAL<br>AQRLAALSTAERREHLAHLIRAEVAAVLGHGDDAAIDRDRAFRDLGFDSMTAVDLRNRLAAVTGVREAATVVFDHPTITRLADHYLE<br>RLVGGSGGGSGNEVTTLENDAAFIENENAYLEKEIARLRKEKAALRNRLAHKKLEHHHHHH |
| MK183 (5)KS1-AT1-KR2-ACP1-SZ3 | MSGDNGMTEEKLRRYLKRTVTELDSVTARLREVEHRAGEPVAVVAMACRLPGGVSTPEEFWELLSEGRDAVAGLPTDRGWDLDSLFH<br>PDPTRSGTAHQRGGGFLTEATAFDPAFFGMSPREALAVDPQQRLMLELSWEVLERAGIPPTSLQASPTGVFVGLIPQEYGPRLAEGG<br>EGVEGYLMTGTTTSVASGRIAYTLGLEGPAISVDTACSSSLVAVHLACQSLRRGESSLAMAGGVTVMPTPGMLVDFSRMNSLAPDGR<br>CKAFSAGANGFGMAEGAGMLLLERLSDARRNGHPVLAVLRGTAVNSDGASNGLSAPNGRAQVRVIQQALAESGLGPADIDAVEAHGT<br>GTRLGDPIEARALFEAYGRDREQPLHLGSVKSNLGHTQAAAGVAGVIKMVLAMRAGTLPRTLHASERSKEIDWSSGAISLLDEPEPW<br>PAGARPRRAGVSSFGISGTNAHAIIEEAPQVVEGERVEAGDVVAPWVLSASSAEGLRAQAARLAAHLREHPGQDPRDIAYSLATGRA<br>ALPHRAAFAPVDESAALRVLDGLATGNADGAAVGTSRAQQRAVFVFPGQGWQWAGMAVDLLDTSPVFAAALRECADALEPHLDFEVI<br>PFLRAEAARREQDAALSTERVDVVQPVMFAVMVSLASMWRAHGVEPAAVIGHSQGEIAAACVAGALSLDDAARVVALRSRVIATMPG<br>NKGMASIAAPAGEVRARIGDRVEIAAVNGPRSVVVAGDSDELDRLVASCTTECIRAKRLAVDYASHSSHVETIRDALHAELGEDFHP<br>LPGFVPFFSTVTGRWTQPDELDAGYWYRNLRRTVRFADAVRALAEQGYRTFLEVSAHPILTAAIEEIGDGSGADLSAIHSLRRGDGS<br>LADFGEALSRAFAAGVAVDWESVHLGTGARRVPLPTYPFQRERVWLLPDRTTPRDELDGWFYRVDWTEVPRSEPAALRGRWLVVVPE<br>GHEEDGWTVEVRSALAEAGAEPEVTRGVGGLVGDCAGVVSLLALEGDGAVQTLVLVRELDAEGIDAPLWTVTFGAVDAGSPVARPDQ<br>AKLWGLGQVASLERGPRWTGLVDLPHMPDPELRGRLTAVLAGSEDQVAVRADAVRARRLSPAHVTATSEYAVPGGTILVTGGTAGLG<br>AEVARWLAGRGAEHLALVSRRGPDTEGVGDLTAELTRLGARVSVHACDVSSREPVRELVHGLIEQGDVVRGVVHAAGLPQQVAINDM<br>DEAAFDEVVAAKAGGAVHLDELCSDAELFLLFSSGAGVWGSARQGAYAAGNAFLDAFARHRRGRGLPATSVAWGLWAAGGMTGDEEA<br>VSFLRERGVRAMPVPRALAALDRVLASGETAVVVTDVDWPAFAESYTAARPRPLLDRIVTTAPSERAGEPETESLRDRLASLPAPER<br>EKALFELVRSHAAAVLGHASAERVPADQAFAELGVDSLSALELRNRLGAATGVRLPTTTVFDHPDVRTLAAHLAAELGGGSGGGSGN<br>EVTTLENDAAFIENENAYLEKEIARLRKEKAALRNRLAHKKLEHHHHHH |

| Construct | Amino acid sequence |
|---|---|
| MK184<br>(5)KS1-<br>AT1-KR1-<br>ACP2-SZ3 | MSGDNGMTEEKLRRYLKRTVTELDSVTARLREVEHRAGEPVAVVAMACRLPGGVSTPEEFWELLSEGRDAVAGLPTDRGWDLDSLFH<br>PDPTRSGTAHQRGGGFLTEATAFDPAFFGMSPREALAVDPQQRLMLELSWEVLERAGIPPTSLQASPTGVFVGLIPQEYGPRLAEGG<br>EGVEGYLMTGTTTSVASGRIAYTLGLEGPAISVDTACSSSLVAVHLACQSLRRGESSLAMAGGVTVMPTPGMLVDFSRMNSLAPDGR<br>CKAFSAGANGFGMAEGAGMLLLERLSDARRNGHPVLAVLRGTAVNSDGASNGLSAPNGRAQVRVIQQALAESGLGPADIDAVEAHGT<br>GTRLGDPIEARALFEAYGRDREQPLHLGSVKSNLGHTQAAAGVAGVIKMVLAMRAGTLPRTLHASERSKEIDWSSGAISLLDEPEPW<br>PAGARPRRAGVSSFGISGTNAHAIIEEAPQVVEGERVEAGDVVAPWVLSASSAEGLRAQAARLAAHLREHPGQDPRDIAYSLATGRA<br>ALPHRAAFAPVDESAALRVLDGLATGNADGAAVGTSRAQQRAVFVFPGQGWQWAGMAVDLLDTSPVFAAALRECADALEPHLDFEVI<br>PFLRAEAARREQDAALSTERVDVVQPVMFAVMVSLASMWRAHGVEPAAVIGHSQGEIAAACVAGALSLDDAARVVALRSRVIATMPG<br>NKGMASIAAPAGEVRARIGDRVEIAAVNGPRSVVVAGDSDELDRLVASCTTECIRAKRLAVDYASHSSHVETIRDALHAELGEDFHP<br>LPGFVPFFSTVTGRWTQPDELDAGYWYRNLRRTVRFADAVRALAEQGYRTFLEVSAHPILTAAIEEIGDGSGADLSAIHSLRRGDGS<br>LADFGEALSRAFAAGVAVDWESVHLGTGARRVPLPTYPFQRERVWLEPKPVARRSTEVDEVSALRYRIEWRPTGAGEPARLDGTWLV<br>AKYAGTADETSTAAREALESAGARVRELVVDARCGRDELAERLRSVGEVAGVLSLLAVDEAEPEEAPLALASLADTLSLVQAMVSAE<br>LGCPLWTVTESAVATGPFERVRNAAHGALWGVGRVIALENPAVWGGLVDVPAGSVAELARHLAAVVSGGAGEDQLALRADGVYGRRW<br>VRAAAPATDDEWKPTGTVLVTGGTGGVGGQIARWLARRGAPHLLLVSRSGPDADGAGELVAELEALGARTTVAACDVTDRESVRELL<br>GGIGDDVPLSAVFHAAATLDDGTVDTLTGERIERASRAKVLGARNLHELTRELDLTAFVLFSSFASAFGAPGLGGYAPGNAYLDGLA<br>QQRRSDGLPATAVAWGTWAGSGMAEGPVADRFRRHGVIEMPPETACRALQNALDRAEVCPIVIDVRWDRFLLAYTAQRPTRLFDEID<br>DARRAAPQAAAEPRVGALAGLPRAERTAELVRLVRTSTATVLGHDDPKAVRATTPFKELGFDSLAAVRLRNLLNAATGLRLPSTLVF<br>DHPNASAVAGFLTSELGGGSGGGSGNEVTTLENDAAFIENENAYLEKEIARLRKEKAALRNRLAHKKLEHHHHHH |
| MK185<br>(5)KS1-<br>AT1-KR1-<br>ACP5-SZ3 | MSGDNGMTEEKLRRYLKRTVTELDSVTARLREVEHRAGEPVAVVAMACRLPGGVSTPEEFWELLSEGRDAVAGLPTDRGWDLDSLFH<br>PDPTRSGTAHQRGGGFLTEATAFDPAFFGMSPREALAVDPQQRLMLELSWEVLERAGIPPTSLQASPTGVFVGLIPQEYGPRLAEGG<br>EGVEGYLMTGTTTSVASGRIAYTLGLEGPAISVDTACSSSLVAVHLACQSLRRGESSLAMAGGVTVMPTPGMLVDFSRMNSLAPDGR<br>CKAFSAGANGFGMAEGAGMLLLERLSDARRNGHPVLAVLRGTAVNSDGASNGLSAPNGRAQVRVIQQALAESGLGPADIDAVEAHGT<br>GTRLGDPIEARALFEAYGRDREQPLHLGSVKSNLGHTQAAAGVAGVIKMVLAMRAGTLPRTLHASERSKEIDWSSGAISLLDEPEPW<br>PAGARPRRAGVSSFGISGTNAHAIIEEAPQVVEGERVEAGDVVAPWVLSASSAEGLRAQAARLAAHLREHPGQDPRDIAYSLATGRA<br>ALPHRAAFAPVDESAALRVLDGLATGNADGAAVGTSRAQQRAVFVFPGQGWQWAGMAVDLLDTSPVFAAALRECADALEPHLDFEVI<br>PFLRAEAARREQDAALSTERVDVVQPVMFAVMVSLASMWRAHGVEPAAVIGHSQGEIAAACVAGALSLDDAARVVALRSRVIATMPG<br>NKGMASIAAPAGEVRARIGDRVEIAAVNGPRSVVVAGDSDELDRLVASCTTECIRAKRLAVDYASHSSHVETIRDALHAELGEDFHP<br>LPGFVPFFSTVTGRWTQPDELDAGYWYRNLRRTVRFADAVRALAEQGYRTFLEVSAHPILTAAIEEIGDGSGADLSAIHSLRRGDGS<br>LADFGEALSRAFAAGVAVDWESVHLGTGARRVPLPTYPFQRERVWLEPKPVARRSTEVDEVSALRYRIEWRPTGAGEPARLDGTWLV<br>AKYAGTADETSTAAREALESAGARVRELVVDARCGRDELAERLRSVGEVAGVLSLLAVDEAEPEEAPLALASLADTLSLVQAMVSAE<br>LGCPLWTVTESAVATGPFERVRNAAHGALWGVGRVIALENPAVWGGLVDVPAGSVAELARHLAAVVSGGAGEDQLALRADGVYGRRW<br>VRAAAPATDDEWKPTGTVLVTGGTGGVGGQIARWLARRGAPHLLLVSRSGPDADGAGELVAELEALGARTTVAACDVTDRESVRELL<br>GGIGDDVPLSAVFHAAATLDDGTVDTLTGERIERASRAKVLGARNLHELTRELDLTAFVLFSSFASAFGAPGLGGYAPGNAYLDGLA<br>QQRRSDGLPATAVAWGTWAGSGMAEGPVADRFRRHGVIEMPPETACRALQNALDRAEVCPIVIDVRWDRFLLAYTAQRPTRLFDEID<br>DARRAAPQAAAEPRVGALAALSTAERREHLAHLIRAEVAAVLGHGDDAAIDRDRAFRDLGFDSMTAVDLRNRLAAVTGVREAATVVF<br>DHPTITRLADHYLERLGGGSGGGSGNEVTTLENDAAFIENENAYLEKEIARLRKEKAALRNRLAHKKLEHHHHHH |
